# Human CSB-deficient iPSCs exhibit impaired DNA damage repair and stress responses following BPDE exposure in an early developmental model

**DOI:** 10.64898/2025.12.02.691678

**Authors:** Alessia Lofrano, Wasco Wruck, Nina Graffmann, James Adjaye

## Abstract

Maintenance of genome integrity is essential for normal human development, particularly during pre-gastrulation stages when rapid proliferation and intense transcriptional activity increase susceptibility to DNA damage. Environmental genotoxins such as benzo[a]pyrene (BaP), a widespread polycyclic aromatic hydrocarbon, and its reactive metabolite benzo[a]pyrene diol epoxide (BPDE) form bulky DNA adducts that interfere with replication and transcription, thereby posing significant risks to embryonic genome stability. To examine how genetic defects in DNA repair influence these effects, we assessed human induced pluripotent stem cells (iPSCs) carrying pathogenic mutations in *ERCC6* (encoding the Cockayne syndrome B, CSB, protein), a key component of transcription-coupled nucleotide excision repair. Pathogenic *ERCC6* mutations result in Cockayne syndrome- a severe neurodevelopmental disorder characterized by growth failure, premature aging, and multisystemic degeneration, thus underscoring the essential developmental functions of CSB. Exposure of healthy and CSB-deficient patient derived iPSCs to BPDE revealed impaired proliferation, persistent accumulation of DNA damage and defective checkpoint activation in CSB-deficient lines. Although the levels of key pluripotency-regulating proteins such as OCT4 and NANOG remained unaltered, we observed altered levels of SOX2 and p-SMAD1/5 signaling thus implying that unrepaired DNA damage can perturb developmental-associated signaling pathways and biological processes. Transcriptomic profiling revealed broad suppression of DNA repair and cell-cycle pathways together with activation of p53-, TNFα-, and MAPK/JNK-mediated stress responses in CSB-deficient lines. Failure to induce anti-oxidant defenses, including SOD2 and IDO, further contributed to oxidative imbalance and incomplete apoptotic clearance. These findings demonstrate that CSB function is essential for coupling DNA repair with transcriptional recovery and redox homeostasis in pluripotent cells. Loss of CSB destabilizes the genome stability under genotoxic stress, providing a mechanistic basis for developmental toxicity of environmental polycyclic aromatic hydrocarbons and underscoring the importance of considering genetic susceptibility in developmental toxicology risk assessment.

## INTRODUCTION

Cockayne syndrome (CS) is a rare autosomal recessive multisystem disorder [1] characterized by growth failure, cataract, photosensitivity, premature aging, and neurodevelopmental delay, with a median life expectancy of approximately 12 years [2]. The clinical spectrum is heterogeneous, ranging from the classical type I form with onset in early childhood to the more severe type II with neonatal manifestations and the milder type III with later onset. The cerebro-oculo-facio-skeletal (COFS) syndrome represents the most extreme variant, often incompatible with long-term survival [3], [4].

CS exhibits genetic heterogeneity, with mutations in distinct genes leading to clinically similar phenotypes. Specifically, CS type A (CSA) results from mutations in the excision repair cross-complementing protein group 8 (*ERCC8)* gene [5], whereas CS type B (CSB) is caused by mutations in *ERCC6* on chromosome 10q11, which encodes the Cockayne syndrome B protein [6].

The CSB protein is a member of the SWI2/SNF2 family of ATP-dependent chromatin remodelers that plays pleiotropic roles in transcription and DNA damage repair. Loss of CSB function impairs the ability of cells to repair transcription-blocking lesions, thus leading to defects in genome stability and transcriptional regulation [7], [8].

CSB plays a central role during the transcription-coupled nucleotide excision repair (TC-NER). CSB recognizes stalled RNA polymerase II at bulky lesions such as UV-induced photoproducts or chemical adducts, and recruits repair factors including CSA, UVSSA, and XPG to promote lesion excision and transcriptional recovery [9],[10],[11]. Importantly, it also supports transcription elongation independent of DNA damage, as such acting as a cofactor of RNA polymerase II and thereby influencing global transcriptional dynamics [6].

Furthermore, CSB has emerged as a regulator of DNA damage response (DDR) to double-strand breaks (DSBs). It facilitates homologous recombination (HR) by recruiting BRCA1, RPA, and RAD51, while loss of CSB favors error-prone non-homologous end joining (NHEJ) via 53BP1–RIF1 foci formation [12]. In addition, CSB deficiency perturbs ATM/CHK2 signaling, leading to premature mitotic entry and genomic instability [13]. Together, these evidence highlight the multifaceted role of CSB in transcriptional homeostasis and DNA repair pathway choice, thus providing a molecular basis for the hypersensitivity of CS cells to genotoxic stress and their contribution to disease pathology and accelerated aging.

Cells are continuously challenged by DNA damage from both endogenous and exogenous genotoxic stressors from environment, diet, and chemicals [13]. Endogenous metabolism alone generates an estimated ∼2,000 – 10,000 lesions per cell per day including oxidative damage and strand breaks, with total damage potentially exceeding 10⁵ lesions daily [14]. The endogenous DNA adducts have been quantified to ∼38,000 adducts per cell [15]. These findings emphasize the urgent need to investigate the molecular mechanisms underlying genotoxic action, particularly in patients with impaired DNA repair capacity, who are prone to accumulations of mutations and chromosomal aberrations.

One environmentally relevant genotoxin is benzo[a]pyrene (BaP), a ubiquitous polycyclic aromatic hydrocarbon (PAH) classified as a group 1 carcinogen according to the International Agency for Research on Cancer (IARC) of the World Health Organization (WHO) [16]. BaP is generated by incomplete combustion of organic materials or fossil fuels and is present in tobacco smoke, vehicle exhaust, charred foods, and industrial emissions [16],[17]. Metabolic activation by cytochrome P450(CYP), generates, amongst others, benzo[a]pyrene-7,8-dihydrodiol-9,10-epoxide (BPDE) [18], which covalently binds DNA, generating bulky adducts that stall RNA polymerase II and activate TC-NER [19]. Persistent BPDE adducts cause replication stress, transcriptional interference, and mutagenesis. Given that TC-NER is initiated by CSB-mediated recognition of stalled transcription complexes, CS-deficient cells are particularly vulnerable to BPDE-induced damage [9]. Despite the well-documented genotoxic potential of BPDE in somatic models, its impact during embryonic development remains underexplored [20].

BaP is capable of crossing the human placenta, exposing the fetus and allowing in situ metabolic activation to BPDE, which forms DNA adducts in both placental tissue and fetal circulation [22]. Similar findings were reported in placentas from smoking mothers, showing BPDE–DNA adducts in both placenta and umbilical cord blood [22]. This is particularly relevant considering that the embryonic period of mammalian development constitutes a critical and highly sensitive phase during which genomic integrity is essential for proper cellular differentiation and organogenesis [23]. Errors in DNA repair mechanisms at this stage can be propagated across proliferating lineages, potentially leading to developmental abnormalities and impaired tissue formation.

Traditionally, developmental toxicity studies have relied on animal models. However, the translational value of these systems is limited by interspecies variability, which often undermines their ability to accurately predict human teratogenic and genotoxic risks. A classic example is thalidomide, which appeared safe in mouse models, yet produced severe malformations in human embryos, thus illustrating the shortcomings of animal-based testing [24].

To overcome these limitations, human pluripotent stem cells have emerged as highly relevant systems for modeling early development and assessing genotoxicity in a human context [23],[24]. Human embryonic stem (ES) cells, derived from the inner cell mass of the blastocyst, display unlimited self-renewal and differentiation potential. Their robust DNA repair mechanisms protect against genomic instability during rapid proliferation [23],[25]. Nevertheless, congenital abnormalities are observed in approximately 3% of live births, with up to 10% of these cases attributed to environmental or chemical exposures [24]. These observations highlight the necessity of elucidating the interplay between DNA repair processes and exogenous stressors during early developmental stages to better understand the etiology of congenital disorders.

Building upon the foundational insights provided by ES cells, human induced pluripotent stem cells (iPSCs) now offer an ethically acceptable and genetically precise alternative for developmental modeling. iPSCs retain the genetic background of the donor, thereby improving the relevance of toxicological and pharmacological assessments and enabling direct investigation of patient-specific molecular mechanisms [26],[27].

In the context of Cockayne syndrome, iPSC-based systems provide a powerful tool to study how deficiencies in transcription-coupled nucleotide excision repair influence the cellular response to environmental genotoxins. These models are particularly well suited for examining the effects of BPDE adducts in a clinically relevant genetic backgrounds, capturing CS-specific vulnerabilities to environmental contaminants. By revealing how BPDE exposure impacts self-renewal capacity and cellular stability in healthy control and CSB iPSC models, this study aims to clarify how defective DNA repair alters developmental homeostasis under genotoxic stress. Ultimately, these insights enhance our understanding of the molecular determinants that safeguard early human development and may provide preventive or therapeutic strategies to mitigate genotoxic risks.

## MATERIALS AND METHODS

### Cell culture

In this study, we utilized a healthy male-derived human induced pluripotent stem cell (hiPSC) line, UM51[28]. Additionally, two hiPSC lines derived from patients with Cockayne Syndrome (CS) were included: IUFi001 and CS789 [29], [30] (Fig. 1A). The IUFi001 line, referred to as CSB_Mild, was derived from dermal fibroblasts of a 3-year-old female diagnosed with Cockayne Syndrome type I. This patient carried compound heterozygous mutations in the *ERCC6* gene: Exon 5 (K377X) and Exon 15 (R857X). The CS789 line, referred to as CSB_Severe, was derived from dermal fibroblasts of an 11-month-old male with cerebro-oculo-facio-skeletal syndrome (COFS), harboring a homozygous *ERCC6* mutation in Exon 10 (R683X). All iPSC lines were cultured on Matrigel-coated plates (Corning) in StemMACS™ iPS-Brew XF medium (Miltenyi). Cells were passaged either as clusters or as single cells. For cluster passaging, cells were detached with room temperature Dulbecco’s Phosphate-Buffered Saline without calcium and magnesium (DPBS; Gibco, Thermo Fisher Scientific), centrifuged at 40xg for 3 minutes and resuspended in StemMACS™ iPS-Brew XF medium. For single-cell passaging, cultures were incubated with pre-warmed Accutase (Life Technologies) for 3 minutes at 37 °C, followed by centrifugation at 190 × g for 3 minutes. Cells were then resuspended in StemMACS™ iPS-Brew XF medium supplemented with 10 µM ROCK inhibitor (Y-27632; Sigma-Aldrich) and maintained with daily medium changes. Cells were cultured at 37 °C, 5% CO_2_ and 5% oxygen Cells were treated at approximately 90% confluency for 24h with BPDE (Benzo[a]pyrene-7,8-dihydrodiol-9,10-epoxide; Santa Cruz Biotechnology) in StemMACS™ iPS-Brew XF medium. BPDE was dissolved in dimethyl sulfoxide (DMSO; Sigma-Aldrich). DMSO was used as a vehicle control in all experiments. Following the 24h treatment period, cells were immediately processed for downstream analyses.

**Figure 1:**
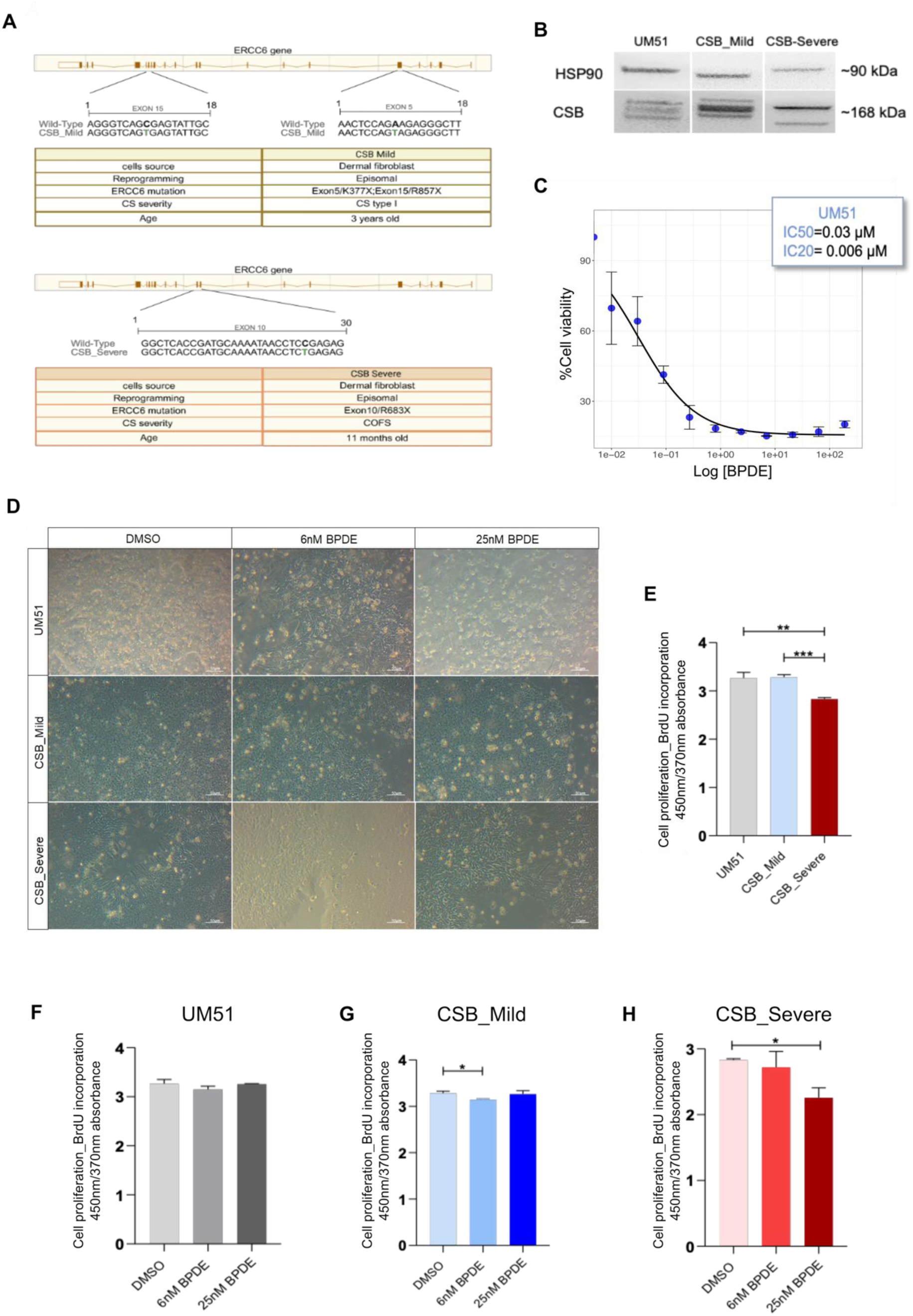
Characterization of ERCC6/CSB mutations and cellular responses to BPDE exposure. **(A)** Schematic representation of the *ERCC6* gene structure and the mutation sites identified in CSB_Mild and CSB_Severe cell lines. Schematic adapted from www.ensembl.org**. (B)** Immunodetection of CSB protein expression in healthy control and CS patient-derived iPSCs. HSP90 was used as a loading control. **(C)** Cell viability dose–response curve of UM51 cells after 24-hour BPDE treatment, assessed by resazurin assay. **(D)** Representative images showing iPSC colony morphology after 24 hours of BPDE exposure at mildly toxic (6 nM) and highly toxic (25 nM) concentrations. Scale bar: 50 µm. **(E)** BrdU incorporation assay measuring proliferation in untreated UM51, CSB_Mild, and CSB_Severe iPSCs. **(F–H)** BrdU incorporation assays assessing proliferation of UM51 (F), CSB_Mild (G), and CSB_Severe (H) iPSCs following 24-hour exposure to mild (6 nM) and toxic (25 nM) BPDE concentrations. (*n* = 3) Statistical significance is indicated as \**p* < 0.05, **p < 0.01, \*\*\**p* < 0.001 and \*\*\*\**p* < 0.0001.

### Cell viability assay

The resazurin assay was employed to determine the optimal concentrations of BPDE to be used for treatment of iPSCs with the aim of establishing the concentrations that induced 20% and 50% (IC20 and IC50) reduction of cell viability. 15.000 UM51 cells/well were seeded in 96-well plates. Upon reaching confluence, cells were exposed to BPDE concentrations ranging from 1nM to 189µM for 24h. 5mg of resazurin (Sigma-Aldrich) were dissolved in 50 mL of sterile DPBS. Twenty hours post-treatment, 20 µL resazurin solution were added to each well, and plates incubated for an additional 4h at 37°C in a 5% CO₂ atmosphere. Fluorescence was measured using a microplate reader (Eppendorf PlateReader AF2200) with excitation at 540nm and emission at 590nm. Fluorescence data were analyzed to generate the dose-response curve. Resazurin measurements of UM51 cells subjected to increasing BPDE doses were normalized according to the manufacturer’s protocol. The normalized values were imported into the R environment [31] and the R packages dr4pl [32] and ggplot2 [33] were used for curve-fitting with a logistic model, plotting and calculation of the IC20 and IC50 values.

### Cell proliferation assay

Cellular proliferation was assessed using the Cell Proliferation ELISA, BrdU (colorimetric) kit (Roche, 1164722900). iPSCs were seeded at a density of 125,000 cells per well in a 96-well plate. Upon reaching approximately 90% confluence, cells were treated with DMSO as a control, 6 nM and 25 nM BPDE, each in 100µL of culture medium, and incubated for 24 hours at 37°C in a 5% CO₂ incubator. Following the 24-hour treatment, 10µL of BrdU labeling solution was added to each well, and cells were incubated for an additional 2h at 37°C in a 5% CO₂ incubator. After incubation, the medium containing the labeling solution was removed, and cells were dried at 60°C for 1h. Subsequently, 200µL of FixDenat solution was added to each well, and cells were incubated at room temperature for 30 minutes. After removing the FixDenat solution, 100µL of Anti-BrdU-POD working solution was added to each well, and cells were incubated at room temperature for 90 minutes. The Anti-BrdU-POD solution was then replaced with 100µL of substrate solution, and cells were incubated at room temperature for up to 30 minutes. Every 5 minutes, absorbance was measured at 370nm with a reference wavelength of 492nm using a microplate reader (Company). The reaction was terminated by adding 25µL of 1 M H₂SO₄ to each well.

### Protein Extraction and Quantification

After 24h of BPDE treatment, iPSCs were harvested using phosphate-buffered saline (PBS) and centrifuged at 500 × g for 3 minutes. Total protein was extracted using RIPA buffer (Sigma-Aldrich) supplemented with protease and phosphatase inhibitors (Roche). The cell suspension was incubated on ice for 1h, followed by centrifugation at 20,000 × g for 10 minutes at 4°C. The supernatant was collected, and protein concentration was determined using the Pierce™ BCA Protein Assay Kit (Thermo Fisher).

### Western blotting

Protein lysates (20–35 µg) were separated by SDS-PAGE using 4–12% gradient gels (Invitrogen, Carlsbad, CA, USA).Proteins were transferred onto a 0.45µm nitrocellulose membrane (Cytiva) in cold conditions for 3h at 300mA. Membranes were blocked with 5% milk (Carl Roth) in Tris-buffered saline with 0.1% Tween-20 (TBS-T) for 1h at room temperature. Primary antibody incubation was performed overnight at 4°C at a dilution of 1:1000. After three washes with TBS-T (5 minutes each), membranes were incubated with secondary antibody (1:2500 dilution) for 1h at room temperature. Antibodies used are listed in Supplementary Table 2. Anti-RPLP0 and Anti-β-ACTIN antibodies were used as housekeeping for normalization. Protein detection was performed using the ECL Western Blotting Detection Reagents (Thermo Fisher) and visualized with a Fusion X imaging system (Peqlab). Band intensity quantification and analysis were performed using FusionCapt Advance software FX7 16.08 (Peqlab).

### Reverse Transcriptase (RT)-qPCR

After 24h of BPDE treatment, total RNA was extracted from iPSCs using the innuPREP RNA Mini Kit 2.0 (Analytik Jena GmbH) following the manufacturer’s instructions for adherent cells. RNA quality and quantity were assessed using a Nanodrop spectrophotometer. Subsequently, 500 ng of RNA was reverse transcribed using the TaqMan Reverse Transcription Kit (Applied Biosystems) to generate complementary DNA (cDNA). RT-qPCR was performed using 50 ng/µL cDNA, Power SYBR™ Green Master Mix (Life Technologies), and primers listed in Supplementary Table1. Amplification was conducted on a VIIA7 Real-Time PCR System (Life Technologies). Each reaction was performed in technical triplicate across three independent biological experiments. Gene expression levels were normalized to the housekeeping gene RPLP0, and fold changes were calculated relative to the DMSO control group using the ΔΔCt method. Statistical comparisons were performed using two-way analysis of variance (ANOVA) with GraphPad Prism v.8.0.2 software (GraphPad Software). A p-value of <0.05 was considered statistically significant.

### Immunocytochemistry

After 24h of BPDE treatment, iPSCs were fixed with 4% paraformaldehyde (PFA) (Polysciences) for 10 minutes at room temperature (RT). Following fixation, cells were washed three times with PBS. For intracellular staining, cells were permeabilized with 0.1% Triton X-100 (Sigma-Aldrich) in PBS for 30 minutes at RT, followed by three 5-minute washes with PBS. To block non-specific binding, cells were incubated with 3% bovine serum albumin (BSA) (Carl Roth) in PBS for 1h at RT. Subsequently, cells were incubated with primary antibodies (diluted 1:400 in 3% BSA) for 2h at RT. For intracellular staining, after primary antibody incubation, cells were washed with 0.05% Tween-20 in PBS and then twice with PBS. Secondary antibodies were diluted 1:400 in 3% BSA supplemented with Hoechst 33258 (Thermo Fisher Scientific) for nuclear counterstaining. After 1h of incubation at RT in the dark, cells were washed as described above. Stained cells were visualized using a Zeiss LSM 700 fluorescence microscope (Carl Zeiss). Individual channel images were processed and merged using ImageJ software (U.S. National Institutes of Health, Bethesda, MD, USA). A list of antibodies used is provided in Supplementary Table 2.

### Stress Array

Protein extraction and quantification were performed as previously described. Cell stress-related proteins were detected using the Human Cell Stress Array Kit (R&D Systems), following the manufacturer’s protocol. 1mL of chemiluminescent detection reagent ECL (Thermo Fisher Scientific) was applied to the membrane, and images were acquired using the Fusion X imaging system (Vilber) after exposure times of 1, 3, and 10 minutes.

### Image analysis of Human Cell Stress Arrays

The images scanned from the hybridized Human Cell Stress Arrays (R&D Systems, Catalog Number ARY018) were imported into the FIJI/ImageJ software [34]. The images were preprocessed and the grid was found semi-automatically as described previously [35]. Spots at the detected grid positions were then quantified via the FIJI Microarray Profile plugin by Bob Dougherty and Wayne Rasband (https://www.optinav.info/MicroArray_Profile.htm, accessed on 06 January 2025).

Array data analysis: The values of the quantified array spots were imported into the R/Bioconductor [36] environment and normalized using values from the reference spots on the array. Up- and down-regulated cytokines for the comparisons between distinct conditions were detected via a detection-p-value in at least one case < 0.05, a p-value from the R limma [37] package <0.05, a ratio >1.2 for up-regulation or a ratio <0.8333 for down-regulation. Hierarchical cluster analysis dendrograms and heatmaps were generated via the heatmap.2 function from The R package gplots [38] using Pearson correlation or Euclidean distance as distance measures.

### Next Generation Sequencing Analysis

Following RNA extraction, 50 ng of total RNA was subjected to 3′ RNA sequencing (3′RNA-Seq) using an Illumina NextSeq 2000 platform. Sequencing was performed at the Core Facility Biomedizinisches Forschungszentrum (BMFZ), Heinrich Heine University Düsseldorf. The HISAT2 software was employed to align Fastq files received from the BMFZ against the GRCh38 genome [39]. HISAT2 was parametrized according to the suggestions from Barruzzo et al.[40]:

hisat2 -p 7 -N 1 -L 20 -i S,1,0.5 -D 25 -R 5 --mp 1,0 --sp 3,0 -x hisatindex/grch38_r109 -U input.fastq.gz -S output.sam

The SAMtools software [41] was used to sort the resulting BAM files. The sorted reads in the BAM files were summarized per gene calling the featurecounts routine from the subread (1.6.1) software package [42] alongside with ENSEMBL annotations from the file Homo_sapiens.GRCh38.109.gtf. Counts per gene were then imported into the R/Bioconductor [43]. Counts were normalized with the voom normalization [44] from the limma package [45] using only expressed genes (counts per million > 1 in at least one sample). Gene expression heatmaps were generated using the heatmap.2 function from the gplots package [46] employing Pearson correlation or Euclidean distance as distance measures. The VennDiagram package [47] was applied for drawing Venn diagrams of expressed genes. Genes were considered expressed when their detection-p-value, calculated as described before [19], when there was more than 1 read per gene. Differentially expressed genes were determined from the intersection set of the venn diagrams using the limma [45] package p < 0.05 and ratios of 1.5 for up- and 0.6667 for down-regulation: Sets of up- and down-regulated or exclusively expressed subsets from the Venn diagrams were subjected to over-representation analysis of gene ontologies (GOs) via the GOstats R package [48] and KEGG (Kyoto Encyclopedia of Genes and Genomes) pathways [49], downloaded in March 2023. The R package ggplot2 [33] was employed to draw the most significant pathways and GO terms (p < 0.05, n < 60) in dotplots.

### Metascape Analysis

Gene sets derived from RNA-Seq data of UM51, CSB_Mild, and CSB_Severe iPSCs treated with DMSO or 25 nM BPDE for 24h were compared using Venn diagrams to identify overlapping and unique differentially expressed genes across experimental groups. These gene subsets were analyzed using Metascape (http://metascape.org), for functional enrichment and pathway analysis. Enriched terms with a minimum of three overlapping genes and an adjusted p-value <0.05 were considered statistically significant.

### TUNEL Assay

Apoptotic cells were detected using the DeadEnd™ Fluorometric TUNEL System (Promega, G3250) in a 24 wells format. 250.000 cells were plated and treated with BPDE for 24h once they reached approximately 90% confluence. Cells were fixed with 4% PFA for 30 minutes at 37°C and subsequently washed twice with PBS for 5 minutes each. Permeabilization was performed at RT for 10 minutes using 0.7% Triton X-100 and 0.2% Tween-20 in PBS. After washing with PBS for 5 minutes, cells were incubated at RT for 10 minutes with 100 µL equilibration buffer per well. For the labeling reaction, 51µL of TdT reaction mixture (45µL equilibration buffer, 5 µL nucleotide mix, and 1 µL recombinant TdT enzyme) was added per well and incubated under appropriate conditions. The reaction was terminated by adding 2× SSC buffer for 15 minutes at RT, followed by three washes with PBS. Cells were then blocked with blocking buffer supplemented with Hoechst stain for 2h at RT. Fluorescently labeled apoptotic cells were visualized using a Zeiss LSM 700 confocal fluorescence microscope (Carl Zeiss) with excitation at 488nm. TUNEL assay quantification was performed using CellSens software by analyzing six images per condition for three biological replicates. TUNEL intensity was normalized to Hoechst staining and expressed relative to the DMSO control. Statistical analyses were conducted using GraphPad Prism 8.

### Statistical Analysis

Statistical analysis was performed using GraphPad Prism version 8.0.2 (GraphPad Software). Data are presented as mean ± standard deviation (SD). Comparisons between groups were conducted using either two-way ANOVA or unpaired Student’s t-test, as appropriate. A p-value of <0.05 was considered statistically significant.

## RESULTS

### 3.1 Impact of ERCC6 Mutations on CSB Protein and iPSC Growth

Our study was performed using two well-established iPSC lines derived from patients with Cockayne syndrome group B (CSB), along with the UM51 iPSC line as a healthy control. The syndromic lines included IUFi001 and CS789. The IUFi001 line was generated from a patient diagnosed with CS type I and carries two mutations in the *ERCC6* gene: p.K377X in exon 5 and p.R857X in exon 15 (Fig. 1A) [29], [30]. The CS789 line was derived from a patient affected by cerebro-oculo-facio-skeletal (COFS) syndrome and harbors a homozygous p.R683X mutation in exon 10 of the *ERCC6* gene (Fig. 1A) [29]. Based on disease severity, IUFi001 was designated as **CSB_Mild**, whereas CS789 was referred to as **CSB_Severe**.

Both mutations introduce premature stop codons, resulting in a predicted truncated CSB proteins. To evaluate the impact of these mutations on CSB protein expression, we performed Western blot analysis (Fig.1B). Multiple bands were detected, which may reflect alternative splicing isoforms, proteolytic cleavage products, or post-translational modifications of the CSB protein. Notably, the mutations in the *ERCC6* gene did not completely abolish CSB signal. Instead, the banding pattern was altered, with a loss of intermediate bands compared to the control, suggesting changes in protein processing and stability.

To determine whether these alterations compromise DNA damage responses, we exposed iPSCs to BPDE, which induces bulky DNA adducts repaired by the transcription-coupled nucleotide excision repair (TC-NER) pathway, where CSB is the initiating factor [6], [50]. A viability assay in UM51 cells established an IC20 of 6 nM for 24h exposure (Fig. 1C). This dose was defined as mildly toxic, whereas 25 nM was selected as a higher challenge concentration. Both doses were subsequently applied to control and CSB-deficient iPSCs.

24h exposure to 6 nM and 25 nM BPDE did not induce noticeable morphological alterations in either the UM51 control or the CSB patient-derived iPSC lines. Cells retained their characteristic round morphology, high nucleus-to-cytoplasm ratio, and ability to form compact colonies (Fig. 1D). However, proliferation was differentially affected. CSB_Mild and CSB_Severe lines displayed reduced colony numbers already at 6 nM BPDE, whereas UM51 exhibited a comparable reduction only at 25 nM. These results corroborate the increased sensitivity of CSB-deficient iPSCs to BPDE.

To asses the effects of the mutations within the *ERCC6* genes on cellular proliferation, a BrdU Cell Proliferation assay was performed. Consistent with previous reports [51], CSB-deficient iPSCs showed inherently reduced proliferation compared to healthy controls (Fig.1E), a phenotype not observed in patient fibroblasts, indicating a specific requirement for CSB in the iPSC state. Among the lines tested, CSB_Severe displayed the strongest growth impairment. Exposure to 25 nM BPDE further suppressed proliferation and DNA synthesis in CSB_Severe cells (Fig. 1H), whereas UM51 and CSB_Mild cells showed no significant change in DNA synthesis even after treatment (Fig. 1F–G).

Together, these findings demonstrate that pathogenic *ERCC6* mutations alter CSB protein processing and expression, resulting in impaired proliferation of patient-derived iPSCs. Moreover, CSB-deficient iPSCs exhibit heightened sensitivity to BPDE, with CSB_Severe cells being particularly vulnerable. These results underscore the essential role of CSB in supporting iPSC growth and survival under both basal conditions and genotoxic stress.

### 3.2 BPDE does not negatively impact pluripotency

Previous studies have shown that iPSCs are particularly vulnerable to ionizing radiation and other genotoxic agents, mainly due to their rapid accumulation of DNA lesions [52]. iPSCs possess robust genome surveillance mechanisms: DNA damage is primarily managed via activation of the p53 pathway, which engages apoptosis, cell-cycle arrest, and senescence programs to prevent the propagation of mutations [53].

Importantly, p53 also plays a dual role in the regulation of pluripotency. While necessary for genomic stability, excessive p53 activity can repress key pluripotency factors, such as NANOG, thereby impairing the self-renewal capacity of these cells [32], [33]. Given this delicate balance, we determined whether our iPSCs were able to preserve and maintain pluripotency following exposure to the genotoxic compound BPDE. To attain this, pluripotency was assessed 24h after low concentration (6 nM) and high concentration (25 nM) BPDE treatment in the healthy control and CSB_patient derived iPSCs lines.

Immunocytochemistry demonstrated that the expression of OCT4, SOX2, and NANOG, key factors maintaining pluripotency, was unaffected compared to untreated controls. (Fig. 2A-C) (Supplementary Fig. 1A-C). Similarly, mRNA levels of these core pluripotency genes remained stable after treatment (Supplementary Fig.1 D-F).

**Figure 2:**
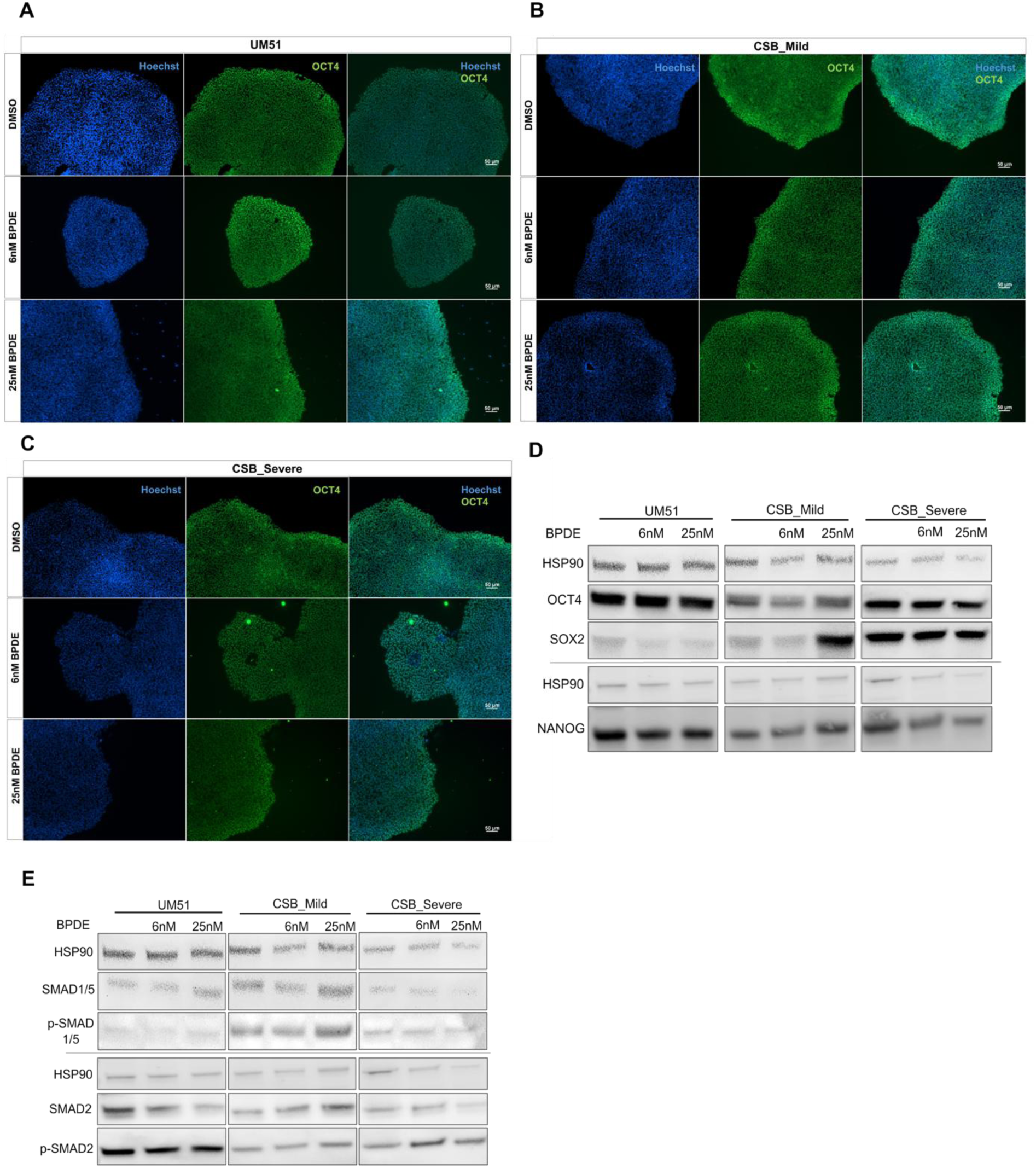
BPDE exposure does not compromise pluripotency in iPSCs. iPSC cultures were treated with 6 nM or 25 nM BPDE for 24 hours prior to fixation or protein/RNA extraction. **(A-C)** Immunofluorescence staining of pluripotency markers OCT4 in UM51 (A), CSB_Mild (B), and CSB_Severe (C) iPSCs following 24-hour exposure to 6 nM or 25 nM BPDE. Scale bar: 50 µm. **(D)** Immunoblot analysis of OCT4, SOX2, and NANOG protein levels in control and CSB iPSCs after BPDE exposure. HSP90 served as a loading control. **(E)** Immunoblot analysis of SMAD1/5 and SMAD2 activation states following 24-hour BPDE treatment in control and CSB iPSCs. HSP90 was used as a loading control.

In contrast, immunoblotting revealed upregulated expression of SOX2 protein in CSB_Mild cells exposed to high dose of BPDE (25 nM), whereas OCT4 and NANOG levels remained unaltered (Fig. 2D). This finding is noteworthy, as SOX2 not only co-operates with OCT4 and NANOG to maintain pluripotency but, when upregulated, can also promote ectodermal differentiation [56], [57].

To investigate this dual role, we analyzed the FGF and SMAD signaling pathway, which is essential for maintaining pluripotency [58]. FGF2 promotes phosphorylation of SMAD2/3, which supports OCT4–SOX2–NANOG transcriptional activity, while simultaneously suppressing the differentiation-inducing factors SMAD1/5/8 [58]. Following BPDE treatment, no significant changes in phosphorylated SMAD2 levels were detected across all cell lines (Fig. 2E). In contrast, CSB_mild iPSCs exposed to 25 nM BPDE displayed an increase in phosphorylated SMAD1/5 (Fig. 2E), consistent with the observed upregulated expression of SOX2.

Taken together, these data imply that BPDE exposure does not compromise the overall pluripotent state of iPSCs. However, in DNA repair–deficient lines such as CSB_mild patient-derived iPSCs, higher BPDE concentrations activate SOX2-dependent pathways, likely in response to accumulated DNA damage, thereby predisposing cells toward differentiation to neuro-ectoderm [59].

### 3.3 Compensatory stress mechanisms preserve Self-Renewal in Healthy but not in CSB-Patient derived iPSCs under BPDE exposure

Although BPDE exposure does not directly impair pluripotency, our findings imply activation of stress-responsive pathways, such as SMAD1/5/8 (Fig. 2E) in CSB_Mild iPSCs, therefore suggesting that iPSCs respond to genotoxic stress not by immediately compromising pluripotency but by fine-tuning expression of lineage-associated factors. To further investigate this adaptive response, we employed a Human Cell stress Array, exposing UM51 and CSB_Severe iPSCs to 25 nM BPDE for 24h to systematically evaluate how stress-associated proteins influence the balance between maintenance of pluripotency and early lineage commitment.

Cluster dendrogram analysis revealed a clear separation between CSB_Severe and UM51 samples, as well as distinct clustering of treated versus untreated conditions (Supplementary Fig. 2A–B). The stress array identified several proteins upregulated in UM51 cells following BPDE exposure (Supplementary Fig. 2C-D), many of which are associated with the maintenance of pluripotency.

Notably, the expression of Disintegrin and Metalloproteinase with Thrombospondin motifs 1 (ADAMTS1), which promotes transforming growth factor beta (TGF-β) induction [60], was upregulated in UM51, suggesting a potential mechanism by which these cells attempt to maintain self-renewal capacity. In contrast, no significant changes in ADAMTS1 expression were detected in CSB_Severe after treatment (Fig.3A-B).

**Figure 3:**
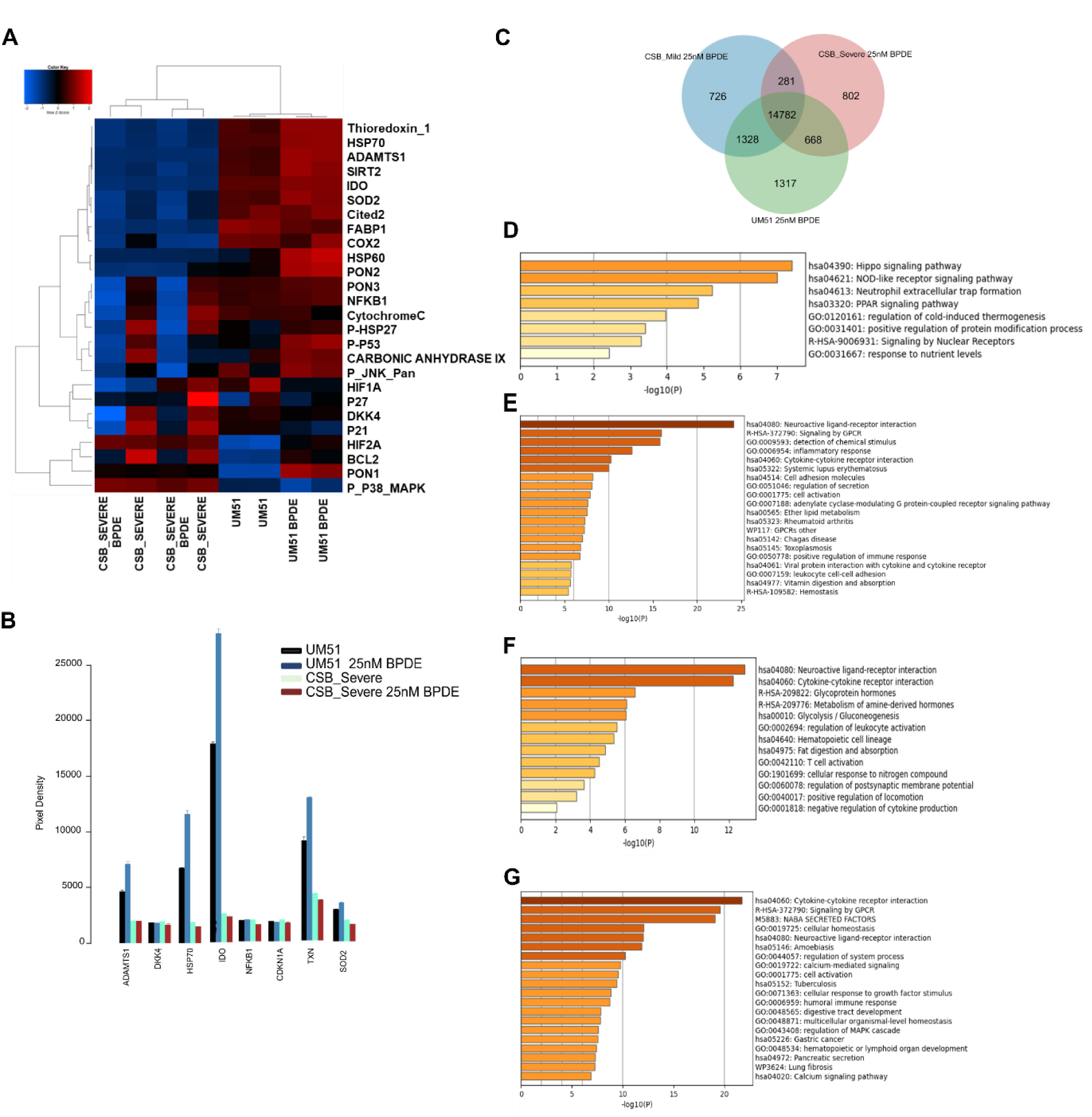
BPDE exposure promotes the pluripotency-associated pathways in healthy control but not in DNA repair-deficient CSB iPSCs. iPSC cultures at approximately 90 % confluency were treated with 25 nM BPDE for 24 hours prior to stress array analysis. **(A)** Pearson’s correlation heatmap showing relative expression levels of key proteins involved in pluripotency maintenance and stress response in control and BPDE-treated iPSCs. **(B)** Bar plot representing the most significantly altered proteins identified by the Human Cell Stress Array analysis. **(C)** Venn diagram depicting genes uniquely or commonly expressed in UM51, CSB_Mild, and CSB_Severe iPSCs after BPDE treatment (detection *p* < 0.05). **(D–G)** Metascape bar graph for viewing top enrichment clusters in: **(D)** genes expressed in common between CSB_Mild and CSB_Severe iPSCs but absent in UM51; **(E)** uniquely expressed in UM51 after 25 nM BPDE exposure**; (F)** uniquely expressed in CSB_Mild after 25 nM BPDE exposure; **(G)** uniquely expressed in CSB_Severe after 25 nM BPDE exposure.

We next examined Indoleamine 2,3-dioxygenase (IDO), a metabolic regulator that supports pluripotency by promoting glycolysis [61]. IDO expression tends to be downregulated in association with the *ERCC6* mutation (Fig. 3A-B). Interestingly, UM51 cells showed a trend toward IDO upregulation following BPDE exposure, suggesting enhanced metabolic activity to preserve cellular homeostasis. In contrast, CSB_Severe cells exhibited unaltered IDO expression (Fig. 3A-B).

Another important secondary regulator of pluripotency is superoxide dismutase 2 (SOD2), which is transcriptionally promoted by OCT4 and NANOG and plays a critical role in limiting reactive oxygen species (ROS) accumulation [62]. Increased ROS levels and oxidative stress induction have previously been reported in CSB patient-derived fibroblasts [63], and this is in line with the observed downregulation of Thioredoxin (TXN) (Fig. 3A-B), a key mediator of cellular antioxidant activity in CSB_Severe compared to UM51. Consistent with this, our results also indicate that SOD2 expression is affected by the *ERCC6* mutation (Fig.3 A_B). Moreover, BPDE exposure led to an upregulated expression of SOD2 in healthy UM51 iPSCs, likely reflecting a compensatory mechanism to sustain pluripotency under stress conditions (Fig. 3A-B). In contrast, CSB_Severe cells exhibited a slight downregulation of SOD2 following BPDE treatment (Fig. 3A-B).

### 3.4 Transcriptomic profiling reveals differential BPDE responses in Healthy and CSB-Deficient iPSCs

To assess the transcriptional consequences of BPDE exposure in iPSCs derived from healthy donors and from CSB patients, bulk RNA sequencing (RNA-seq) on three independent iPSC lines under untreated conditions and following 24h exposure to 25 nM BPDE was performed. Differential gene expression analysis identified 281 genes that were commonly expressed in CSB patient-derived iPSCs but absent in the control line UM51, 802 uniquely expressed in CSB_severe, 726 uniquely expressed in CSB_Mild and 1317 uniquely expressed in UM51 (Fig. 3C).

Functional enrichment analysis of these transcripts was performed using Metascape. The results revealed a significant over-representation of the Hippo signaling pathway (Fig. 3D) in common between the CSB_patient derived iPSC lines after BPDE treatment. Activation of Hippo signaling has been associated with sustained cellular stress responses, including transcriptional stress, persistent DNA lesions, chromatin remodeling, and elevated oxidative stress, all of which are characteristic features of CSB-deficient cells following genotoxic exposure [64]. The enrichment of this pathway therefore supports the hypothesis that CSB-deficient iPSCs experience prolonged stress signaling in response to BPDE-induced DNA damage.

Metascape analysis also unveiled upregulation of the NOD-like receptor (NOD) signaling pathway in CSB patient derived iPSCs following BPDE exposure, which is typically activated by intracellular danger-associated molecular patterns (DAMPs) and cellular stress [65]. NOD pathway activation can trigger downstream NF-κB and MAPK cascades, consistent with the enrichment of MAPK regulatory genes detected among the CSB-severe transcripts (Fig. 3G). This finding suggests that persistent DNA damage and stress-induced cytoplasmic signaling may contribute towards the activation of inflammation-associated pathways in CSB iPSCs.

However, the Metascape analysis of the CSB_mild iPSC line revealed an enrichment of pathways involved in metabolic homeostasis and glycolytic regulation, indicating that these cells may preferentially engage adaptive metabolic programs to restore cellular energy balance following genotoxic insult (Fig. 3F).

To better assess the transcriptional response to BPDE, we performed pairwise comparisons between treated and control conditions for each cell line (p < 0.05). A Venn diagram identified unique and shared expressed genes across conditions (Supplementary Fig. 3A). In healthy iPSCs, 283 genes were uniquely expressed after BPDE exposure, primarily associated with developmental and immune processes (Supplementary Fig. 3B-C), whereas 314 genes were uniquely expressed in untreated controls. Of the 17,267 genes shared between conditions, differential expression analysis followed by KEGG and GO enrichment revealed no major DNA damage involved pathway enrichment indicative of a strong BPDE-specific response (Supplementary Fig. 3E-F), consistent with the notion that iPSCs mount a rapid DNA damage response to preserve genomic stability.

The transcriptional response of CSB patient-derived iPSCs following BPDE exposure revealed more pronounced effects. CSB_Mild iPSCs exposed to 25 nM BPDE for 24h displayed 465 genes uniquely expressed upon treatment and 511 genes expressed only in untreated controls (Fig. 4A). Among the 16,741 commonly expressed genes, KEGG analysis revealed upregulation of drug metabolism pathways, including Cytochrome P450 enzymes involved in BPDE detoxification, and pathways associated with cell death (Fig. 4B). Metabolic pathways were consistently downregulated in CSB_Mild treated iPSCs, in agreement with the stress array results (Fig. 3A-4C). Cell cycle–related pathways were also significantly downregulated, thus suggesting a protective mechanism to preserve genomic integrity (Fig. 4C–F).

**Figure 4:**
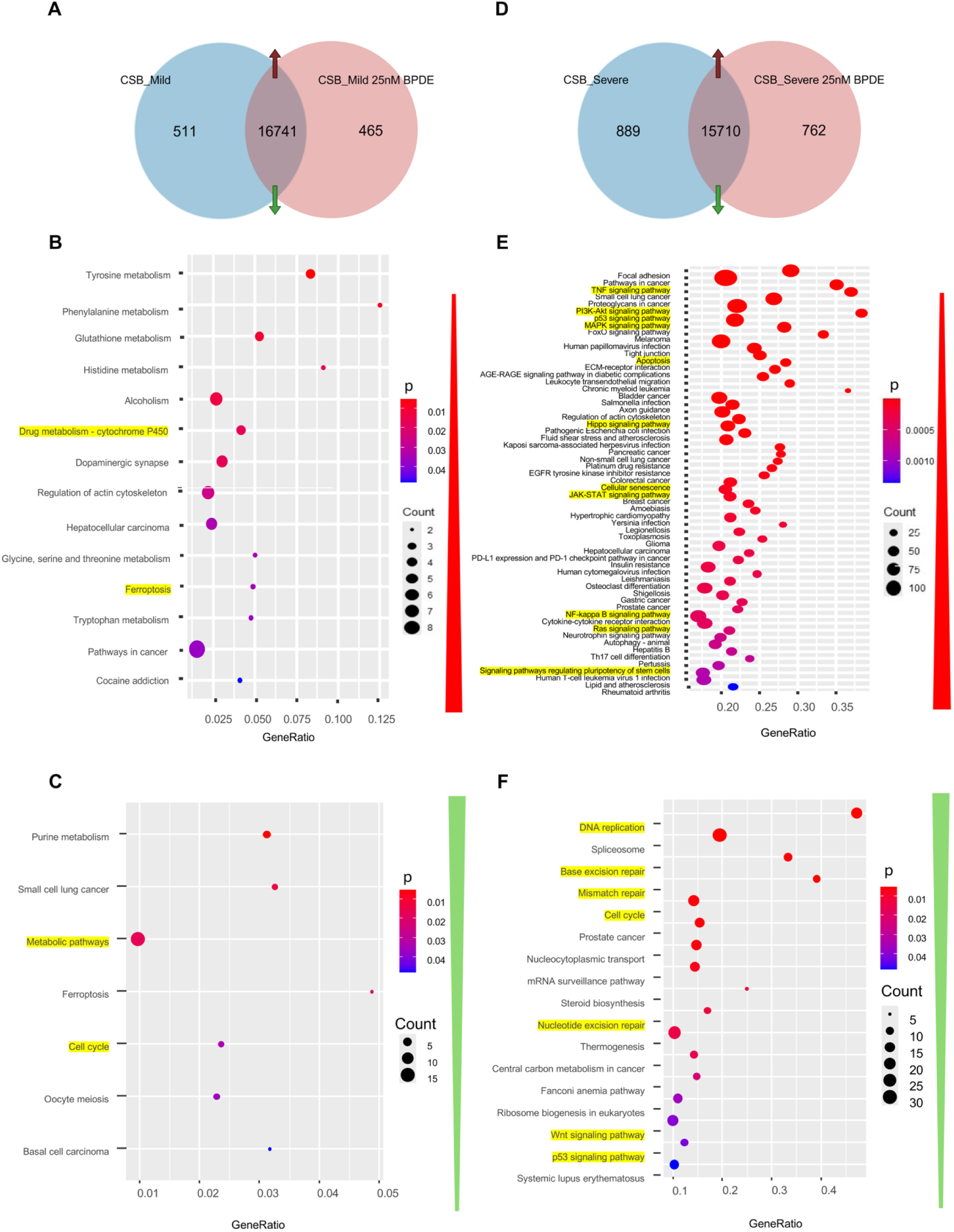
Differential gene expression and pathway enrichment in CSB-deficient iPSCs following BPDE exposure: **(A)** Venn diagram illustrating the distribution of genes uniquely expressed under control conditions (511), exclusively expressed following 24-hour exposure to 25 nM BPDE (465), and commonly expressed (16,741) in CSB_Mild iPSCs (*p* < 0.05, detection threshold). **(B)** Dot plot showing the top upregulated KEGG pathways among the 16,741 differentially expressed genes (DEGs) in CSB_Mild iPSCs exposed to 25 nM BPDE compared with untreated controls. **(C)** Dot plot showing the top downregulated KEGG pathways among the 16,741 DEGs in CSB_Mild iPSCs following BPDE exposure. **(D)** Venn diagram representing genes uniquely expressed under control conditions (689), exclusively expressed after 24-hour exposure to 25 nM BPDE (762), and commonly expressed (15,710) in CSB_Severe iPSCs (*p* < 0.05, detection threshold). **(E)** Dot plot summarizing the top upregulated KEGG pathways among DEGs in CSB_Severe iPSCs following 25 nM BPDE exposure relative to untreated controls. **(F)** Dot plot summarizing the top downregulated KEGG pathways among DEGs in CSB_Severe iPSCs following 25 nM BPDE exposure relative to untreated controls.

CSB_Severe cells displayed the most profound transcriptional dysregulation (Fig. 4D). DNA damage–associated pathways, including DNA replication, base excision repair, and mismatch repair, were suppressed, a pattern not observed in CSB_Mild cells (Fig. 4F). This indicates that CSB_Severe cells fail to efficiently resolve BPDE-induced DNA lesions, reflecting a collapse of the DNA damage response (DDR) on a background already compromised by loss of functional CSB protein. Consequently, these cells are likely to accumulate mutations, undergo apoptosis, and activate additional stress response pathways.

Pathway enrichment analysis confirmed strong upregulation of TNF-α, PI3K–Akt, p53, and MAPK signaling cascades. TNF-α mediates inflammatory and stress responses, while p53 activation controls cell cycle arrest, DNA repair, apoptosis, and pluripotency regulation. In CSB_Severe cells, pro-apoptotic p53 target genes (e.g., *BBC3, BCL2L1, CASP3, CCND1, CD82, CDK6, CDKN1A, CDKN2A, DDB2, FAS, PERP*) were upregulated (Supplementary Table 3), whereas p53-associated genes involved in cell cycle arrest and DNA repair (*CASP9, CCNB1, CCNE1, CHEK2*) were downregulated, indicating failure to restore genomic stability [66], [67], [68], [69] (Supplementary Table 4). KEGG and stress array analyses further confirmed that CSB_Severe cells fail to activate compensatory metabolic programs required to maintain pluripotency under genotoxic stress, suggesting susceptibility to differentiation defects or loss of self-renewal not yet apparent after 24 h.

In parallel, robust induction of MAPK/JNK and PI3K–AKT (Fig. 4E) pathways highlight a dual response: MAPK/JNK promotes apoptosis via modulation of p53 activity, whereas PI3K–AKT provides pro-survival signals. This balance between pro-apoptotic and pro-survival signaling underscores the complex integration of stress responses in CSB-deficient iPSCs.

### 3.5 CSB iPSCs following BPDE treatment show impaired DNA Damage Response when compared to Healthy control

The embryotoxicity-associated properties of BPDE and its downstream DNA damage dynamics remain poorly understood in human, with only limited studies addressing this issue [21], [22]. To fill this gap, we investigated BPDE-induced DNA damage accumulation and DDR dynamics in two systems-healthy iPSCs and CSB-deficient iPSCs, in which the central DDR protein CSB is defective. Previous studies in somatic cells have shown that CSB deficiency impairs lesion removal [70].

In pluripotent cells, UV-induced 6–4 photoproducts are typically removed within ∼2h, while cyclobutane pyrimidine dimers are resolved more slowly (6–24h), with overall repair kinetics faster than in somatic cells [61]. Based on these kinetics and given the use of a DNA repair–deficient model, we selected 24h as the time point to assess BPDE effects.

To determine whether BPDE activates the nucleotide excision repair (NER) pathway, we analyzed the transcriptional response of two key lesion-recognition factors [71]: Xeroderma Pigmentosum Group C (XPC) and DNA Damage-Binding Protein 2 (DDB2). *XPC* expression remained unchanged across all lines (Supplementary Fig. 5A). A ∼2-fold induction of *DDB2* was detected in CSB_Mild cells after 25 nM BPDE treatment, whereas no significant changes were observed in CSB_Severe or UM51 cells (Supplementary Fig. 5B).

Given this limited transcriptional activation of NER components, we next examined whether BPDE exposure results in the accumulation of double-strand breaks (DSBs). Although BPDE does not directly induce DSBs, impaired NER can generate replication-associated DSBs through stalled forks and uncoupled lagging-strand synthesis [72]. To assess this, we examined the effect of BPDE on γH2AX levels.

After 24h, UM51 cells showed no significant increase in γH2AX, consistent with efficient and fast repair of transient breaks (Fig. 5A) (Fig. D-E). By contrast, both CSB lines exhibited dose-dependent γH2AX accumulation (Fig. 5B–D).

**Figure 5:**
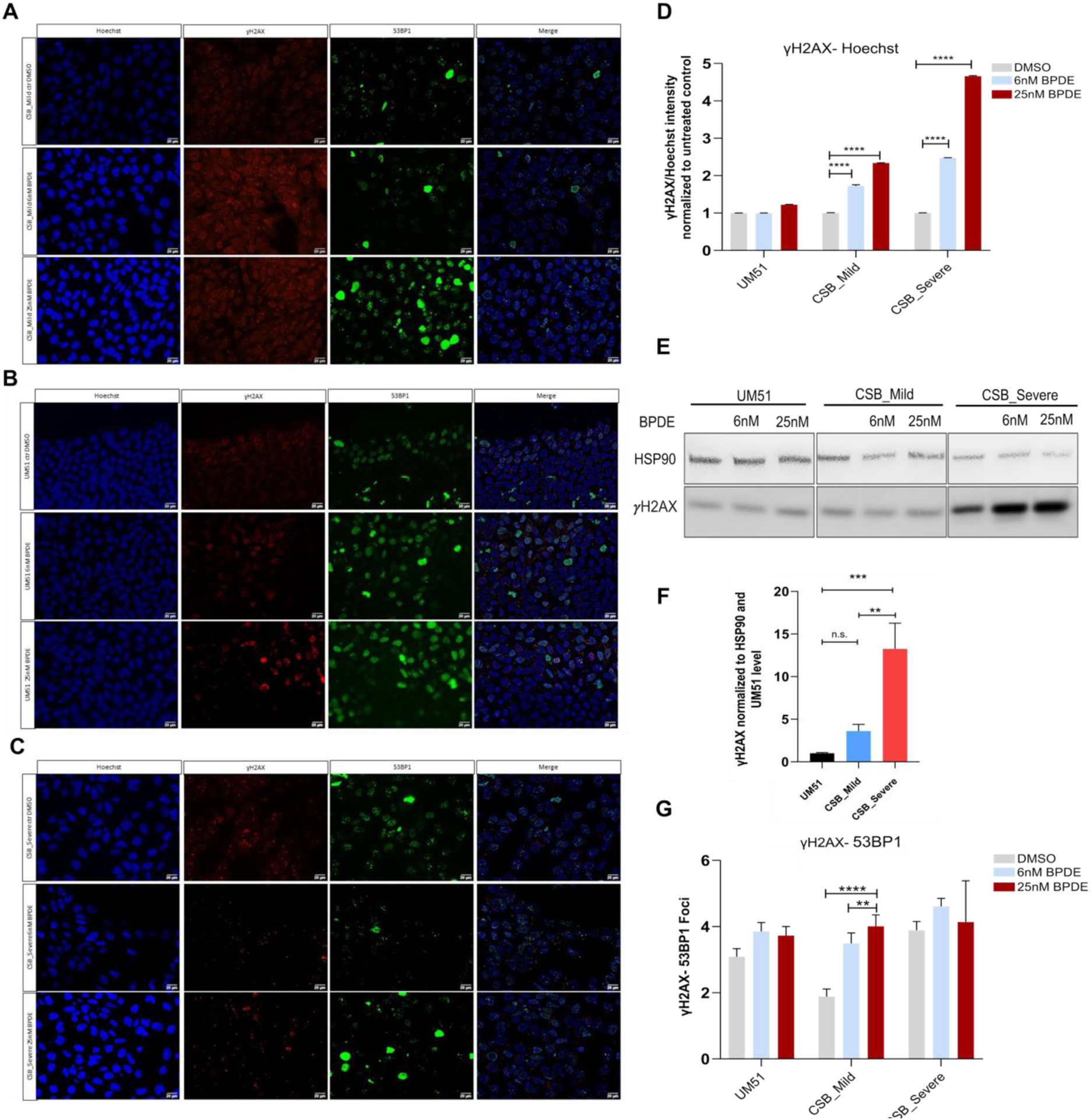
Impaired DNA damage response in CSB-deficient iPSCs following BPDE treatment compared with healthy controls. iPSC cultures at approximately 90 % confluency were treated with 6 nM or 25 nM BPDE for 24 hours prior to fixation or protein/RNA extraction. **(A-C)** Immunofluorescence staining of γH2AX (red)–53BP1 (green) after 24-hour BPDE treatment in UM51 (A), CSB_Mild (B), and CSB_Severe (C) iPSCs. **(D)** Quantification of γH2AX was normalized to Hoechst nuclear staining. Five images per biological replicate (n = 3) were analyzed. **(E)**Immunoblot analysis of γH2AX protein levels in UM51, CSB_Mild, and CSB_Severe iPSCs after BPDE exposure. **(F)** Densitometric quantification of γH2AX normalized to HSP90 and expressed relative to healthy control UM51 (n = 3). **(G)** Immunostaining and quantification of γH2AX–53BP1 foci formation following BPDE exposure. Foci were manually counted from five images per biological replicate (n = 3) and normalized to untreated controls. Statistical significance is indicated as \**p* < 0.05, **p < 0.01, \*\*\**p* < 0.001 and \*\*\*\**p* < 0.0001.

γH2AX, a central DDR marker, recruits repair-associated proteins such as 53BP1 to damage sites, coordinating either repair or apoptosis. To further characterize DDR activation, we evaluated γH2AX–53BP1 foci formation after 24h 6 nM and 25 nM BPDE treatment. Healthy iPSCs showed no foci induction (Fig. 5A and 5G), whereas CSB_Mild cells displayed a robust, dose-dependent increase (Fig. 5B and 5G). CSB_Severe cells, however, failed to form foci, suggesting collapse of DDR signaling (Fig. 5C and 5G).

These results indicate that healthy iPSCs efficiently recognize and repair BPDE lesions, such that by 24h most damage is repaired. In contrast, CSB deficiency delays DDR kinetics, leading to persistent γH2AX and 53BP1 activation (Fig. 5G).

Importantly, CSB patient-derived iPSCs also displayed elevated baseline DNA damage and DSBs in the absence of BPDE (Fig. 5E-F). This intrinsic instability is consistent with defective repair of endogenous replication-associated lesions. Thus, both CSB_Mild and CSB_Severe iPSCs appear to exist under chronic stress, characterized by spontaneous DNA damage accumulation (Fig. 5E-F).

### 3.6 DNA repair-deficient iPSCs undergo apoptosis following BPDE exposure

When DNA damage in iPSCs cannot be effectively repaired, cells typically engage protective mechanisms such as cell-cycle arrest or apoptosis to maintain genomic stability [59].

To further elucidate the fate of BPDE-treated iPSCs under conditions of compromised DDR and accumulation of DNA lesions, we analyzed pathways involved in apoptotic and cell cycle regulation. Gene-set enrichment analysis revealed a significant upregulation of cell death–associated processes in both CSB-derived iPSC lines (Fig. 4).

In UM51 iPSCs, BPDE exposure induced phosphorylation of ATM (p-ATM) and CHK2 (p-CHK2) (Fig. 6A) and upregulation of *CHK1* transcription (Fig. 6D). However p53 levels remained largely unchanged at both the mRNA and protein levels (Fig. 7C-D). This response likely reflects activation of a checkpoint-mediated cell-cycle arrest, allowing time for DNA repair therefore suggesting a rapid and functional activation of the ATM–CHK2–p53 axis in response to BPDE treatment. Despite the absence of significant upregulation of γH2AX (Fig. 5D) and p53 after 24h of treatment (Fig. 7D), BAX is upregulated, both at mRNA and protein levels in UM51 (Fig. 7B-D). Notably, cleaved caspase-3 levels were unchanged (Fig. 7D), however the proportion of TUNEL-positive cells increased (Fig. 7E), indicating DNA fragmentation and priming of apoptotic cell death. These findings suggest that the genotoxic stress induced heterogeneous responses within the UM51-iPSC population: most cells likely repaired mild or transient DNA lesions efficiently in the 24h, whereas a subset of cells harboring persistent DNA damage underwent apoptosis.

**Figure 6:**
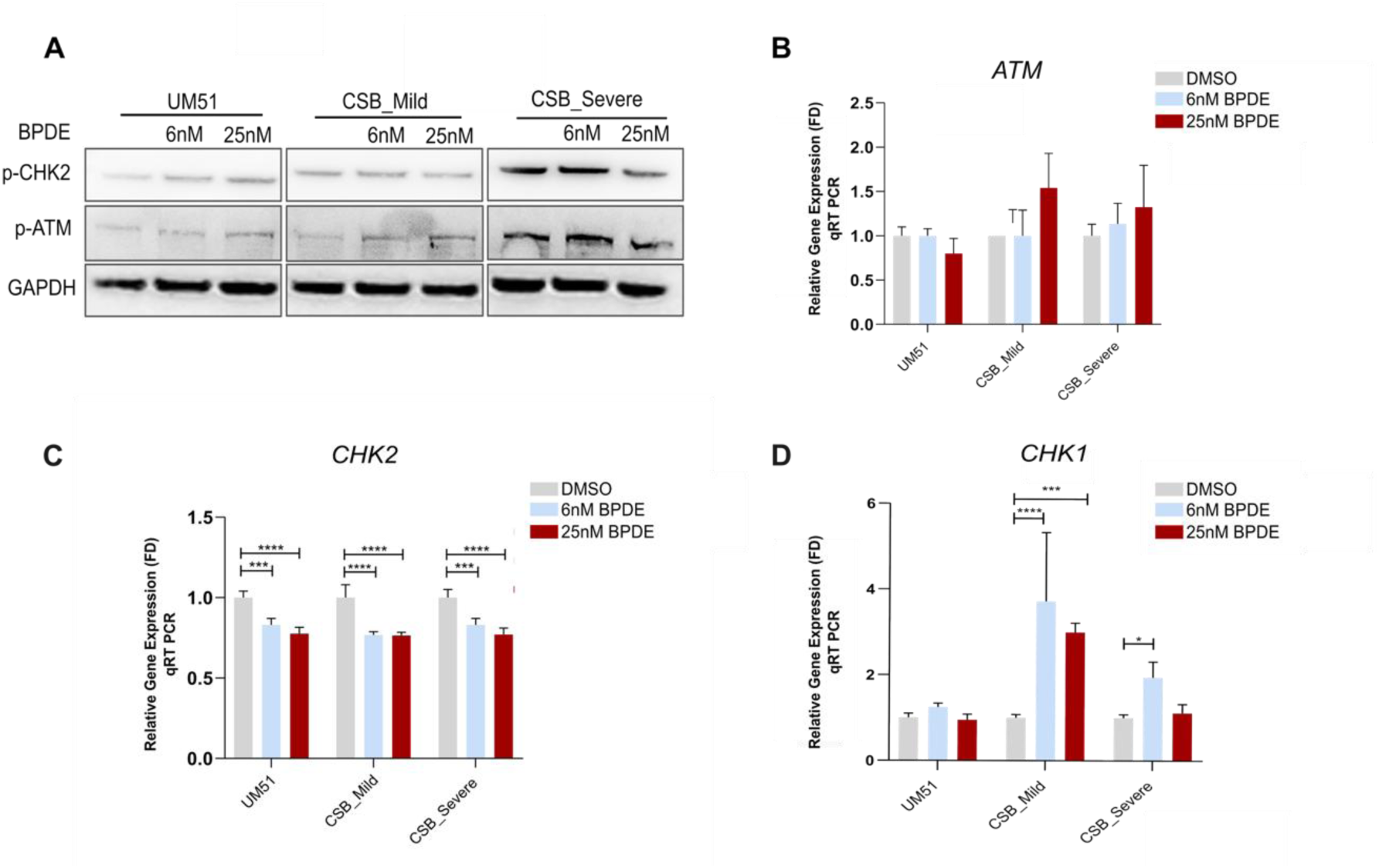
BPDE induces cell cycle arrest in healthy iPSCs but not in CSB-deficient lines. iPSC cultures at approximately 90 % confluency were treated with BPDE for 24 hours prior to fixation or protein/RNA extraction. **(A)** Immunoblot analysis of phosphorylated (activated) CHK2 (Thr68) and phosphorylated ATM protein levels in UM51 and CSB patient–derived iPSCs following BPDE exposure. GAPDH served as a loading control. **(B–D)** Quantitative RT-PCR analysis of *ATM* (B), *CHK2* (C), and *CHK1* (D) transcript levels after BPDE treatment. Expression values were normalized to *RPLP0* and are presented as mean ± 95 % confidence interval (*n* = 3). Statistical significance is indicated as \**p* < 0.05, **p < 0.01, \*\*\**p* < 0.001 and \*\*\*\**p* < 0.0001; colors in the graphs correspond to significance levels as indicated in the figure legend.

**Figure 7:**
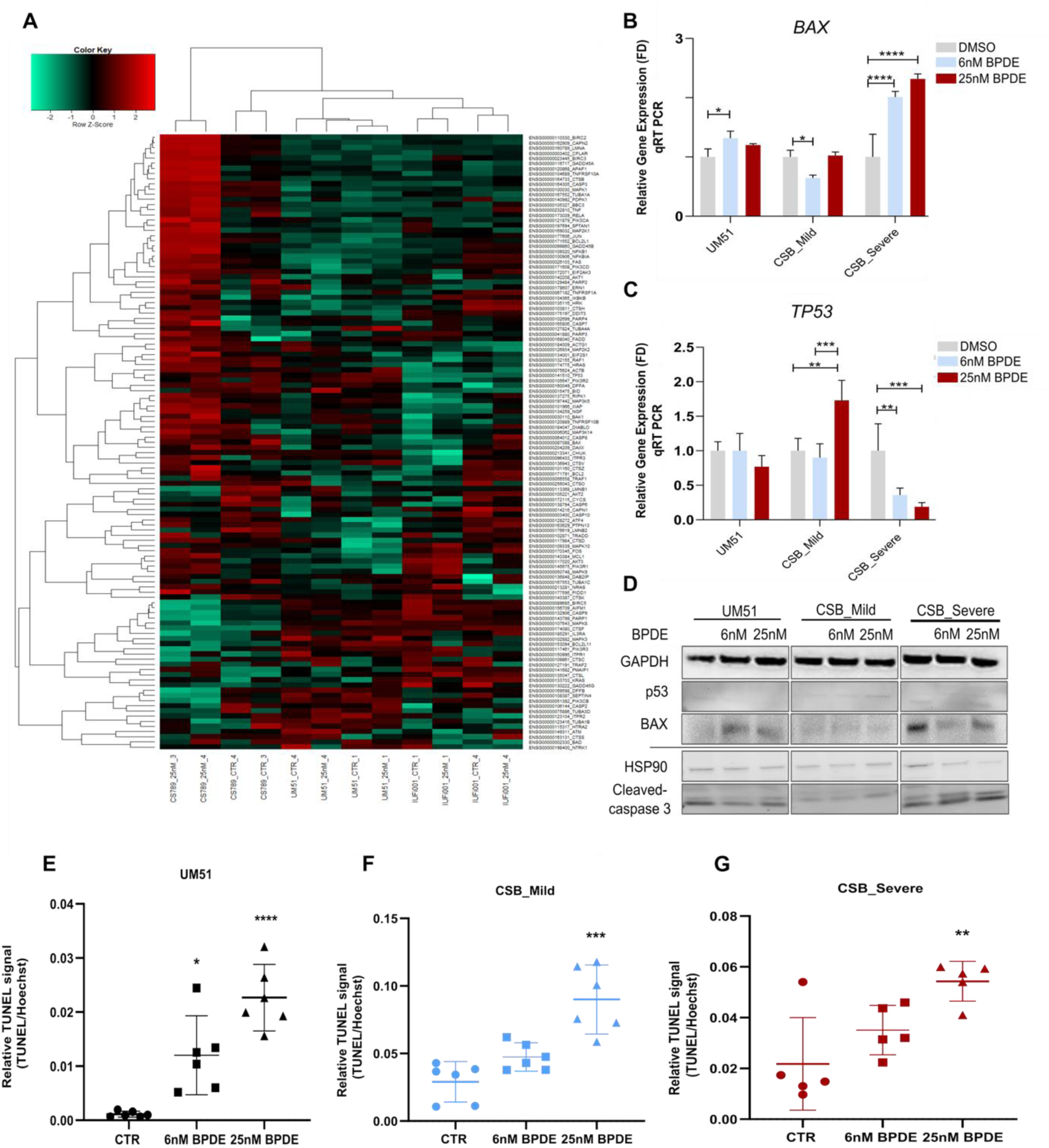
DNA repair–deficient iPSCs undergo apoptosis following BPDE exposure. iPSC cultures at approximately 90 % confluency were treated with 6 nM or 25 nM BPDE for 24 hours prior to fixation or protein/RNA extraction. **(A)** Pearson’s correlation heatmap showing relative expression of apoptosis-associated genes in healthy (UM51) and CSB-deficient (CSB_Mild, CSB_Severe) iPSC lines after 24-hour exposure to 25 nM BPDE. **(B-C)** Quantitative RT-PCR analysis of *BAX* (B) and *TP53* (C) transcript levels in UM51, CSB_Mild, and CSB_Severe iPSCs following BPDE exposure. Expression values were normalized to *RPLP0* and are presented as mean ± 95 % confidence interval (*n* = 3; *p* < 0.05, \**p* < 0.01, \*\**p* < 0.001. **(D)** Immunoblot analysis of p53, BAX and Cleaved-Caspase3 protein levels in UM51, CSB_Mild, and CSB_Severe iPSCs, with GAPDH and HSP90 used as a loading control. **(E–G)** Quantification of TUNEL/Hoechst fluorescence intensity ratio in UM51 (E), CSB_Mild (F), and CSB_Severe (G) iPSCs after 24-hour exposure to 6 nM or 25 nM BPDE compared to the untreated ctr. Six images per biological replicate (*n* = 3) were analyzed. Data represent mean ± 95 % confidence interval; statistical significance is indicated as \**p* < 0.05, **p < 0.01, \*\*\**p* < 0.001 and \*\*\*\**p* < 0.0001.

In contrast, CSB Severe iPSCs exhibited a complete absence of DDR activation 24 hours post-BPDE exposure (Fig. 5G), which correlated with decreased p-ATM and p-CHK2 levels (Fig. 6A) and reduced *TP53* expression (Fig. 7C). Although *BAX* transcription was markedly increased, (Fig. 7B) protein levels were downregulated at low BPDE doses and only slightly upregulated at higher doses (Fig. 7D). These cells also displayed pronounced cleaved caspase-3 (Fig. 7D) activation and positive TUNEL staining after 24h 25 nM BPDE exposure (Fig. 7G), indicative of progression to late-stage apoptosis. Notably, untreated CSB_Severe cells exhibited elevated basal BAX expression (Fig. 7D), suggesting that chronic endogenous DNA damage primes these cells for apoptosis, a phenomenon previously described in CSB-deficient contexts [50]

CSB_Mild iPSCs exhibited a delayed DDR induction following BPDE exposure, as evidenced by increased p-ATM levels (Fig. 6A) and slightly decreased p-CHK2 levels (Fig. 6A-C) along with a 2.5-fold upregulation of *TP53* expression (Fig. 7C). BAX and Cleaved-caspase3 protein levels remained stable (Fig. 7D). Notably, cells showed accumulation of positive TUNEL staining after treatment (Fig.7F), indicating the presence of DNA strand breaks and the initiation of apoptosis. Transcriptional upregulation of *CHK1* in both CSB patient-derived lines (Fig. 6D) further suggests activation of ATR-mediated replication stress signaling, which may partially compensate for impaired ATM–CHK2–p53 signaling. Together, these data imply that CSB deficiency leads to delayed and partially compromised DDR activation, with replication stress responses being invoked to maintain genome integrity, but ultimately resulting in early apoptotic events, as reflected by TUNEL positivity. These observations are in line with previous reports indicating that CHK2 is dispensable for BPDE-induced S-phase checkpoint activation but becomes critical under replication fork stalling, and that CSB deficiency compromises ATM–CHK2 signaling, thereby increasing genomic instability and apoptotic susceptibility [73], [74].

## DISCUSSION

Maintenance of genome integrity is a central requirement for correct human development, particularly during pre-gastrulation stages when rapid cell division and extensive transcriptional activity increase susceptibility to DNA damage. During this critical window, the fidelity of DNA repair and transcriptional recovery pathways determines the stability of the developing genome and the proper execution of developmental programs.

Human are continuously exposed to a complex mixture of environmental genotoxins originating from dietary sources, air pollution, contaminated water, and consumer products. Quantitative exposure assessments indicate that adults typically inhale or ingest microgram-to-nanogram amounts of recognized genotoxins each day, including polycyclic aromatic hydrocarbons (PAHs) such BaP. The most toxic BaP metabolite, BPDE, forms bulky covalent adducts with DNA. If not efficiently repaired, these can lead to mutations, chromosomal instability and long-term epigenetic alterations.

CS represents a proto-typical disorder arising from defects in the transcription-coupled nucleotide excision repair (TC-NER) – pathway, which is essential for resolving transcription-blocking DNA lesions. Mutations in *ERCC6*, which encodes the CSB protein, compromise the removal of bulky DNA adducts and hinder the resumption of transcription following damage. This failure in lesion repair and transcriptional recovery underlies the profound neurodevelopmental and systemic abnormalities characteristic of the disease.

In the present study, we employed human iPSCs as a model to investigate the impact of genotoxic stress during early developmental stages in healthy and disease conditions. Using a CSB-deficient iPSC-based model, we examined the transcriptional and DNA damage responses induced by BPDE-a well established potent embryotoxic and mutagenic environmental agent. This approach allowed us to explore how exposure to environmentally relevant genotoxins influences genome maintenance and transcriptional regulation in pluripotent cells (a surrogate in vitro cell representing the inner cells mass of the blastocyts) during embryonic development.

Our study establishes that pathogenic *ERCC6* mutations alter CSB protein, resulting in impaired proliferation and heightened genotoxic sensitivity in patient-derived iPSCs. Both mild and severe CSB-deficient lines exhibited reduced colony formation under basal conditions, which was exacerbated following BPDE exposure. This vulnerability was especially pronounced in CSB_Severe cells, consistent with the clinical severity of the underlying mutations.

BPDE exposure did not compromise pluripotency per se, as OCT4, SOX2, and NANOG expression remained largely unaltered across all lines. However, subtle transcriptional perturbations, most notably the selective upregulation of SOX2 and p-SMAD1/5 protein expression in CSB_Mild after BPDE treatment and CSB_Severe even in control condition suggest early ectodermal lineage priming in response to unresolved DNA damage. These findings imply that while iPSCs can preserve their pluripotent transcriptional circuitry under moderate genotoxic stress, CSB-deficient cells exhibit altered signaling dynamics that may predispose them to differentiation bias or developmental defects when stress persists.

Through stress-protein profiling, we identified compensatory mechanisms that sustain pluripotency under BPDE exposure in healthy iPSCs but fail in CSB-deficient lines. UM51 control cells activated ADAMTS1, IDO and SOD2-factors that support pluripotency by regulating metabolic and antioxidant activity. In contrast, CSB_Severe iPSCs displayed a lack of induction of these protective factors and downregulation of redox regulators such as TXN and SOD2, suggesting an inability to mount an adequate antioxidant response. This deficiency likely amplifies the accumulation of reactive oxygen species (ROS) and oxidative DNA damage, thus leading to compromised genomic stability and self-renewal.

Transcriptomic analysis reinforced these observations. BPDE exposure elicited limited transcriptional changes in healthy iPSCs, reflecting efficient DNA repair and transcriptional recovery. In contrast, CSB-deficient iPSCs showed extensive transcriptional reprogramming, characterized by suppression of DNA repair and cell-cycle pathways, activation of p53, TNF-α and MAPK/JNK-dependent stress associated cascades. The concurrent activation of apoptotic and survival signaling pathways imply a finely poised but ultimately unsustainable stress response in the absence of functional CSB. In particular in CSB_Severe cells, the collapse of DNA replication, base excision repair and mismatch repair pathways underscores the global failure of genome maintenance mechanisms following BPDE exposure.

Consistent with this transcriptomic profile, functional assays confirmed persistent accumulation of DNA damage and impaired DDR activation in CSB-deficient iPSCs. Both CSB_Mild and CSB_Severe lines accumulated γH2AX foci upon BPDE exposure, but only the milder line retained partial 53BP1 recruitment, thus indicative of residual repair capacity. In contrast, the absence of foci formation and DDR signaling in CSB_Severe iPSCs rafter BPDE exposure reflects a collapse of damage recognition and checkpoint activation. Moreover CSB_severe iPSCs also displayed elevated baseline γH2AX levels, highlighting chronic endogenous DNA damage accumulation even without exogenous genotoxin exposure. Together, these findings imply that CSB-deficient iPSCs exist in a state of constitutive genotoxic stress that compromises the ability to tolerate external stress and induce a collapse of DNA damage response with adverse implications for both proliferation and developmental potential.

Our findings indicate that healthy iPSCs exhibit robust checkpoint activation and selectively undergo apoptosis in cells with unrepairable DNA damage, therefore preventing the accumulation of genomic lesions and reflecting a transient and efficient repair of DNA damage. In contrast, CSB_Severe iPSCs show impaired activation of the ATM–p53–CHK2 axis, leading to defective cell-cycle regulation. Despite BAX induction and increased TUNEL positivity at 25 nM BPDE, these cells fail to mount a fully coordinated apoptotic response, resulting in the accumulation of extensive DNA damage. Overall, these results delineate a mechanistic continuum from partial checkpoint activation and compensatory stress responses in CSB_Mild cells to complete collapse of the DNA damage response in CSB_Severe cells. The differential severity of these phenotypes mirrors the clinical spectrum of Cockayne syndrome and highlights the predictive value of iPSC-based models for genotype–phenotype correlations.

Our findings have broader implications beyond Cockayne syndrome. They highlight the essential role of CSB in orchestrating DNA repair, transcriptional regulation and stress adaptation in pluripotent cells-functions that are fundamental for healthy early embryonic development. Given the known ability of Benzo[a]pyrene and its metabolites to cross the placental barrier, our results provide a mechanistic rationale for the developmental toxicity associated with environmental polycyclic aromatic hydrocarbons. The heightened sensitivity of CSB-deficient iPSCs to BPDE underscores the potential risk of environmental genotoxins in individuals with impaired DNA repair capacity and emphasizes the importance of integrating genetic susceptibility into risk assessment frameworks.

In conclusion, our study reveals that CSB plays a pivotal role in preserving genomic stability and self-renewal capacity under genotoxic stress in pluripotent cells. Loss of CSB function compromises DNA repair disrupts stress-responsive signaling, and predisposes cells to apoptosis and differentiation defects. By leveraging patient-derived iPSCs, we have established a human-relevant model that captures the developmental vulnerability associated with transcription-coupled repair deficiency and provides a powerful platform for investigating the interplay between genetic and environmental factors in early human development. Future studies employing differentiated germ layer-specific cell types may further elucidate tissue-specific mechanisms of DNA damage intolerance and identify therapeutic strategies to mitigate genotoxic risks during embryogenesis.

## Institutional Review Board Statement

The study was conducted according to the guidelines by the Ethics Committee of the medical faculty of Heinrich-Heine University, Germany (protocol code: 5704).

## Author Contribution

A.L. designed and performed experiments, processed, and analysed the data, wrote and edited the manuscript. W.W. performed the bioinformatic analysis, data curation, helped with the figures, wrote the bioinformatic section in the methods and materials, and edited the manuscript. N.G. edited the manuscript, acquired funding and supervised the study. J.A. conceptualized and designed the work, edited the manuscript, acquired funding, and supervised the study. All authors have read and agree to the submitted version of the manuscript.

## REFERENCES

[1] M. A. Nance and S. A. Berry, ‘Cockayne syndrome: Review of 140 cases’, Am. J. Med. Genet., vol. 42, no. 1, pp. 68–84, Jan. 1992, doi: 10.1002/ajmg.1320420115.

[2] T. Y. Abdel Ghaffar, E. S. Elsobky, and S. M. Elsayed, ‘Cholestasis in patients with Cockayne syndrome and suggested modified criteria for clinical diagnosis’, Orphanet J Rare Dis, vol. 6, no. 1, p. 13, Dec. 2011, doi: 10.1186/1750-1172-6-13.

[3] V. Laugel et al., ‘Mutation update for the *CSB* / *ERCC6* and *CSA* / *ERCC8* genes involved in Cockayne syndrome’, Hum. Mutat., vol. 31, no. 2, pp. 113–126, Feb. 2010, doi: 10.1002/humu.21154.

[4] V. Natale, ‘A comprehensive description of the severity groups in Cockayne syndrome’, American J of Med Genetics Pt A, vol. 155, no. 5, pp. 1081–1095, May 2011, doi: 10.1002/ajmg.a.33933.

[5] A. Chebly, S. Corbani, J. Abou Ghoch, C. Mehawej, A. Megarbane, and E. Chouery, ‘First molecular study in Lebanese patients with Cockayne syndrome and report of a novel mutation in ERCC8 gene’, BMC Med Genet, vol. 19, no. 1, p. 161, Dec. 2018, doi: 10.1186/s12881-018-0677-7.

[6] ‘troelstra, C., et al. (1992). ERCC6, a member of a subfamily of putative helicases, is involved in Cockayne’s syndrome and preferential repair of active genes. Cell, 71(6), 939–953. doi:10.1016/0092-8674(92)90390-6.’.

[7] A. van Gool, J. de Wit, W. Vermeulen, D. Bootsma, and J. H. J. Hoeijmakers, ‘ERCCG, a Member of a Subfamily of Putative Helicases, Is Involved in Cockaynes Syndrome and Preferential Repair of Active Genes’.

[8] J. C. Newman, A. D. Bailey, and A. M. Weiner, ‘Cockayne syndrome group B protein (CSB) plays a general role in chromatin maintenance and remodeling’, Proc. Natl. Acad. Sci. U.S.A., vol. 103, no. 25, pp. 9613–9618, Jun. 2006, doi: 10.1073/pnas.0510909103.

[9] P. C. Hanawalt and G. Spivak, ‘Transcription-coupled DNA repair: two decades of progress and surprises’, Nat Rev Mol Cell Biol, vol. 9, no. 12, pp. 958–970, Dec. 2008, doi: 10.1038/nrm2549.

[10] J. A. Marteijn, H. Lans, W. Vermeulen, and J. H. J. Hoeijmakers, ‘Understanding nucleotide excision repair and its roles in cancer and ageing’, Nat Rev Mol Cell Biol, vol. 15, no. 7, pp. 465–481, Jul. 2014, doi: 10.1038/nrm3822.

[11] V. Laugel, ‘Cockayne syndrome: The expanding clinical and mutational spectrum’, Mechanisms of Ageing and Development, vol. 134, no. 5–6, pp. 161–170, May 2013, doi: 10.1016/j.mad.2013.02.006.

[12] R. J. Lake, A. Geyko, G. Hemashettar, Y. Zhao, and H.-Y. Fan, ‘UV-Induced Association of the CSB Remodeling Protein with Chromatin Requires ATP-Dependent Relief of N-Terminal Autorepression’, Molecular Cell, vol. 37, no. 2, pp. 235–246, Jan. 2010, doi: 10.1016/j.molcel.2009.10.027.

[13] ‘Lindahl, T. “Instability and decay of the primary structure of DNA.” Nature vol. 362,6422 (1993): 709–15. doi:10.1038/362709a0’.

[14] ‘Barnes, Deborah E, and Tomas Lindahl. “Repair and genetic consequences of endogenous DNA base damage in mammalian cells.” Annual review of genetics vol. 38 (2004): 445–76. doi:10.1146/annurev.genet.38.072902.092448’.

[15] J. A. Swenberg et al., ‘Endogenous versus Exogenous DNA Adducts: Their Role in Carcinogenesis, Epidemiology, and Risk Assessment’, Toxicological Sciences, vol. 120, no. Supplement 1, pp. S130–S145, Mar. 2011, doi: 10.1093/toxsci/kfq371.

[16] D. N. Das and N. Ravi, ‘Influences of polycyclic aromatic hydrocarbon on the epigenome toxicity and its applicability in human health risk assessment’, Environmental Research, vol. 213, p. 113677, Oct. 2022, doi: 10.1016/j.envres.2022.113677.

[17] ‘Hattemer-Frey, H A, and C C Travis. “Benzo-a-pyrene: environmental partitioning and human exposure.” Toxicology and industrial health vol. 7,3 (1991): 141–57. doi:10.1177/074823379100700303’.

[18] K. Shiizaki, M. Kawanishi, and T. Yagi, ‘Modulation of benzo[a]pyrene–DNA adduct formation by CYP1 inducer and inhibitor’, Genes and Environ, vol. 39, no. 1, p. 14, Dec. 2017, doi: 10.1186/s41021-017-0076-x.

[19] W. Li et al., ‘Human genome-wide repair map of DNA damage caused by the cigarette smoke carcinogen benzo[a]pyrene’, Proc. Natl. Acad. Sci. U.S.A., vol. 114, no. 26, pp. 6752–6757, Jun. 2017, doi: 10.1073/pnas.1706021114.

[20] G. Talaska, M. Jaeger, R. Reilman, T. Collins, and D. Warshawsky, ‘Chronic, topical exposure to benzo[a]pyrene induces relatively high steady-state levels of DNA adducts in target tissues and alters kinetics of adduct loss.’, Proc. Natl. Acad. Sci. U.S.A., vol. 93, no. 15, pp. 7789–7793, Jul. 1996, doi: 10.1073/pnas.93.15.7789.

[21] V. Karttunen, P. Myllynen, G. Prochazka, O. Pelkonen, D. Segerbäck, and K. Vähäkangas, ‘Placental transfer and DNA binding of benzo(a)pyrene in human placental perfusion’, Toxicology Letters, vol. 197, no. 2, pp. 75–81, Aug. 2010, doi: 10.1016/j.toxlet.2010.04.028.

[22] J. Arnould, P. Verhoest, V. Bach, J. Libert, and J. Belegaud, ‘Detection of benzo[a]pyrene-DNA adducts in human placenta and umbilical cord blood’, Hum Exp Toxicol, vol. 16, no. 12, pp. 716–721, Dec. 1997, doi: 10.1177/096032719701601204.

[23] M. N. Shahbazi and V. Pasque, ‘Early human development and stem cell-based human embryo models’, Cell Stem Cell, vol. 31, no. 10, pp. 1398–1418, Oct. 2024, doi: 10.1016/j.stem.2024.09.002.

[24] S. Tandon and S. Jyoti, ‘Embryonic stem cells: An alternative approach to developmental toxicity testing’, J Pharm Bioall Sci, vol. 4, no. 2, p. 96, 2012, doi: 10.4103/0975-7406.94808.

[25] T. M. Kim, V. I. Rebel, and P. Hasty, ‘Defining a genotoxic profile with mouse embryonic stem cells’, Exp Biol Med (Maywood), vol. 238, no. 3, pp. 285–293, Mar. 2013, doi: 10.1177/1535370213480700.

[26] K. Takahashi and S. Yamanaka, ‘Induction of Pluripotent Stem Cells from Mouse Embryonic and Adult Fibroblast Cultures by Defined Factors’, Cell, vol. 126, no. 4, pp. 663–676, Aug. 2006, doi: 10.1016/j.cell.2006.07.024.

[27] ‘Alessandro Prigione, Beatrix Fauler, Rudi Lurz, Hans Lehrach, James Adjaye, The Senescence-Related Mitochondrial/Oxidative Stress Pathway is Repressed in Human Induced Pluripotent Stem Cells, Stem Cells, Volume 28, Issue 4, April 2010, Pages 721–733, 10.1002/stem.404’.

[28] M. Bohndorf, A. Ncube, L.-S. Spitzhorn, J. Enczmann, W. Wruck, and J. Adjaye, ‘Derivation and characterization of integration-free iPSC line ISRM-UM51 derived from SIX2-positive renal cells isolated from urine of an African male expressing the CYP2D6 *4/*17 variant which confers intermediate drug metabolizing activity’, Stem Cell Research, vol. 25, pp. 18–21, Dec. 2017, doi: 10.1016/j.scr.2017.10.004.

[29] L.-P. Szepanowski et al., ‘Cockayne Syndrome Patient iPSC-Derived Brain Organoids and Neurospheres Show Early Transcriptional Dysregulation of Biological Processes Associated with Brain Development and Metabolism’, Cells, vol. 13, no. 7, p. 591, Mar. 2024, doi: 10.3390/cells13070591.

[30] S. Martins, I. Hacheney, N. Teichweyde, B. Hildebrandt, J. Krutmann, and A. Rossi, ‘Generation of an induced pluripotent stem cell line (IUFi001) from a Cockayne syndrome patient carrying a mutation in the ERCC6 gene’, Stem Cell Research, vol. 55, p. 102456, Aug. 2021, doi: 10.1016/j.scr.2021.102456.

[31] R. Ihaka and R. Gentleman, ‘R: A Language for Data Analysis and Graphics’, Journal of Computational and Graphical Statistics, vol. 5, no. 3, pp. 299–314, Sep. 1996, doi: 10.2307/1390807.

[32] H. An, J. Landis T., A. Bailey G., J. Marron S., and D. Dittmer P., ‘dr4pl: A Stable Convergence Algorithm for the 4 Parameter Logistic Model’, The R Journal, vol. 11, no. 2, p. 171, 2019, doi: 10.32614/RJ-2019-003.

[33] H. Wickham, Ggplot2: elegant graphics for data analysis. in Use R! New York: Springer, 2009.

[34] ‘Schneider, Caroline A., Wayne S. Rasband, and Kevin W. Eliceiri. 2012. “NIH Image to ImageJ: 25 Years of Image Analysis.” Nature Methods 9 (7): 671–75. 10.1038/nmeth.2089.’.

[35] ‘Wruck, Wasco, Vincent Boima, Lars Erichsen, et al. 2022. “Urine-Based Detection of Biomarkers Indicative of Chronic Kidney Disease in a Patient Cohort from Ghana.” Journal of Personalized Medicine 13 (1): 38. 10.3390/jpm13010038’.

[36] ‘Gentleman, Robert C, Vincent J Carey, Douglas M Bates, et al. 2004. “Bioconductor: Open Software Development for Computational Biology and Bioinformatics.” Genome Biology 5 (10): R80. 10.1186/gb-2004-5-10-r80’.

[37] ‘Smyth, Gordon K. 2004. “Linear Models and Empirical Bayes Methods for Assessing Differential Expression in Microarray Experiments.” Statistical Applications in Genetics and Molecular Biology 3 (1). 10.2202/1544-6115.1027.’.

[38] ‘Warnes, Gregory R., Ben Bolker, Lodewijk Bonebakker, et al. 2015. Gplots: Various R Programming Tools for Plotting Data. http://CRAN.R-project.org/package=gplots.’.

[39] D. Kim, B. Langmead, and S. L. Salzberg, ‘HISAT: a fast spliced aligner with low memory requirements’, Nature Methods, vol. 12, no. 4, pp. 357–360, Mar. 2015, doi: 10.1038/nmeth.3317.

[40] G. Baruzzo, K. E. Hayer, E. J. Kim, B. Di Camillo, G. A. FitzGerald, and G. R. Grant, ‘Simulation-based comprehensive benchmarking of RNA-seq aligners’, Nature Methods, vol. 14, no. 2, pp. 135–139, Feb. 2017, doi: 10.1038/nmeth.4106.

[41] H. Li et al., ‘The Sequence Alignment/Map format and SAMtools’, Bioinformatics, vol. 25, no. 16, pp. 2078–2079, Aug. 2009, doi: 10.1093/bioinformatics/btp352.

[42] Y. Liao, G. K. Smyth, and W. Shi, ‘featureCounts: an efficient general purpose program for assigning sequence reads to genomic features’, Bioinformatics, vol. 30, no. 7, pp. 923–930, Apr. 2014, doi: 10.1093/bioinformatics/btt656.

[43] R. C. Gentleman et al., ‘Bioconductor: open software development for computational biology and bioinformatics’, Genome Biol., vol. 5, no. 10, p. R80, 2004, doi: 10.1186/gb-2004-5-10-r80.

[44] C. W. Law, Y. Chen, W. Shi, and G. K. Smyth, ‘voom: precision weights unlock linear model analysis tools for RNA-seq read counts’, Genome Biology, vol. 15, no. 2, p. R29, 2014, doi: 10.1186/gb-2014-15-2-r29.

[45] G. K. Smyth, ‘Linear Models and Empirical Bayes Methods for Assessing Differential Expression in Microarray Experiments’, Statistical Applications in Genetics and Molecular Biology, vol. 3, no. 1, 2004, doi: 10.2202/1544-6115.1027.

[46] G. R. Warnes et al., gplots: Various R Programming Tools for Plotting Data. 2015. [Online]. Available: http://CRAN.R-project.org/package=gplots

[47] H. Chen and P. C. Boutros, ‘VennDiagram: a package for the generation of highly-customizable Venn and Euler diagrams in R’, BMC Bioinformatics, vol. 12, no. 1, p. 35, 2011, doi: 10.1186/1471-2105-12-35.

[48] S. Falcon and R. Gentleman, ‘Using GOstats to test gene lists for GO term association’, Bioinformatics, vol. 23, no. 2, pp. 257–258, Jan. 2007, doi: 10.1093/bioinformatics/btl567.

[49] M. Kanehisa, M. Furumichi, M. Tanabe, Y. Sato, and K. Morishima, ‘KEGG: new perspectives on genomes, pathways, diseases and drugs’, Nucleic Acids Res., vol. 45, no. D1, pp. D353–D361, Jan. 2017, doi: 10.1093/nar/gkw1092.

[50] N. L. Batenburg, E. L. Thompson, E. A. Hendrickson, and X. Zhu, ‘Cockayne syndrome group B protein regulates DNA double - strand break repair and checkpoint activation’, The EMBO Journal, vol. 34, no. 10, pp. 1399–1416, May 2015, doi: 10.15252/embj.201490041.

[51] L. N. D. S. Andrade, J. L. Nathanson, G. W. Yeo, C. F. M. Menck, and A. R. Muotri, ‘Evidence for premature aging due to oxidative stress in iPSCs from Cockayne syndrome’, Human Molecular Genetics, vol. 21, no. 17, pp. 3825–3834, Sep. 2012, doi: 10.1093/hmg/dds211.

[52] M. Zhang, C. Yang, H. Liu, and Y. Sun, ‘Induced Pluripotent Stem Cells Are Sensitive to DNA Damage’, Genomics, Proteomics & Bioinformatics, vol. 11, no. 5, pp. 320–326, Oct. 2013, doi: 10.1016/j.gpb.2013.09.006.

[53] H. Wang, M. Guo, H. Wei, and Y. Chen, ‘Targeting p53 pathways: mechanisms, structures and advances in therapy’, Sig Transduct Target Ther, vol. 8, no. 1, p. 92, Mar. 2023, doi: 10.1038/s41392-023-01347-1.

[54] X. Fu, S. Wu, B. Li, Y. Xu, and J. Liu, ‘Functions of p53 in pluripotent stem cells’, Protein Cell, vol. 11, no. 1, pp. 71–78, Jan. 2020, doi: 10.1007/s13238-019-00665-x.

[55] T. Lin and Y. Lin, ‘p53 switches off pluripotency on differentiation’, Stem Cell Res Ther, vol. 8, no. 1, p. 44, Dec. 2017, doi: 10.1186/s13287-017-0498-1.

[56] K. Adachi, H. Suemori, S. Yasuda, N. Nakatsuji, and E. Kawase, ‘Role of *SOX2* in maintaining pluripotency of human embryonic stem cells’, Genes to Cells, vol. 15, no. 5, pp. 455–470, May 2010, doi: 10.1111/j.1365-2443.2010.01400.x.

[57] H. Y. Zhou et al., ‘A *Sox2* distal enhancer cluster regulates embryonic stem cell differentiation potential’, Genes Dev., vol. 28, no. 24, pp. 2699–2711, Dec. 2014, doi: 10.1101/gad.248526.114.

[58] ‘Boris Greber, Hans Lehrach, James Adjaye, Fibroblast Growth Factor 2 Modulates Transforming Growth Factor β Signaling in Mouse Embryonic Fibroblasts and Human ESCs (hESCs) to Support hESC Self-Renewal, Stem Cells, Volume 25, Issue 2, February 2007, Pages 455–464, 10.1634/stemcells.2006-0476’.

[59] W. Chen et al., ‘DNA repair proteins cooperate with SOX2 in regulating the transition of human embryonic stem cells to neural progenitor cells’, Biochemical and Biophysical Research Communications, vol. 586, pp. 163–170, Jan. 2022, doi: 10.1016/j.bbrc.2021.11.060.

[60] ‘Bourd-Boittin, Katia et al. “Protease profiling of liver fibrosis reveals the ADAM metallopeptidase with thrombospondin type 1 motif, 1 as a central activator of transforming growth factor beta.” Hepatology (Baltimore, Md.) vol. 54,6 (2011): 2173–84. doi:10.1002/hep.24598’.

[61] ‘Liu, Xin, et al. “IDO1 Maintains Pluripotency of Primed Human Embryonic Stem Cells by Promoting Glycolysis.” Stem cells (Dayton, Ohio) vol. 37,9 (2019): 1158–1165. doi:10.1002/stem.3044’.

[62] C. Solari et al., ‘Manganese Superoxide Dismutase Gene Expression Is Induced by Nanog and Oct4, Essential Pluripotent Stem Cells’ Transcription Factors’, PLoS ONE, vol. 10, no. 12, p. e0144336, Dec. 2015, doi: 10.1371/journal.pone.0144336.

[63] B. Pascucci et al., ‘An altered redox balance mediates the hypersensitivity of Cockayne syndrome primary fibroblasts to oxidative stress’, Aging Cell, vol. 11, no. 3, pp. 520–529, Jun. 2012, doi: 10.1111/j.1474-9726.2012.00815.x.

[64] ‘Pefani, Dafni E, and Eric O’Neill. “Hippo pathway and protection of genome stability in response to DNA damage.” The FEBS journal vol. 283,8 (2016): 1392–403. doi:10.1111/febs.13604’.

[65] P. J. Murray, ‘NOD proteins: an intracellular pathogen-recognition system or signal transduction modifiers?’, Current Opinion in Immunology, vol. 17, no. 4, pp. 352–358, Aug. 2005, doi: 10.1016/j.coi.2005.05.006.

[66] ‘Bennett M, Macdonald K, Chan SW, Luzio JP, Simari R, Weissberg P. Cell surface trafficking of Fas: a rapid mechanism of p53-mediated apoptosis. Science. 1998 Oct 9;282(5387):290–3. doi: 10.1126/science.282.5387.290. PMID: 9765154.’.

[67] ‘Hussar, P. Apoptosis Regulators Bcl-2 and Caspase-3. Encyclopedia 2022, 2, 1624–1636. 10.3390/encyclopedia2040111’.

[68] ‘Martha L. Slattery, Lila E. Mullany, Roger K. Wolff, Lori C. Sakoda, Wade S. Samowitz, Jennifer S. Herrick, The p53-signaling pathway and colorectal cancer: Interactions between downstream p53 target genes and miRNAs, Genomics, Volume 111, Issue 4, 2019, Pages 762–771, ISSN 0888-7543, 10.1016/j.ygeno.2018.05.006.’.

[69] ‘Attardi LD, Reczek EE, Cosmas C, Demicco EG, McCurrach ME, Lowe SW, Jacks T. PERP, an apoptosis-associated target of p53, is a novel member of the PMP-22/gas3 family. Genes Dev. 2000 Mar 15;14(6):704–18. Erratum in: Genes Dev 2000 Jul 15;14(14):1835. PMID: 10733530; PMCID: PMC316461.’.

[70] L. Z. Luo et al., ‘Zheng H, Wang X, Warren AJ, Legerski RJ, Nairn RS, Hamilton JW, et al. Nucleotide excision repair- and polymerase eta-mediated error-prone removal of mitomycin C interstrand crosslinks. Mol Cell Biol. 2003;23:754–761. doi: 10.1128/MCB.23.2.754-761.2003’, PLoS ONE, vol. 7, no. 3, p. e30541, Mar. 2012, doi: 10.1371/journal.pone.0030541.

[71] A. Ray, K. Milum, A. Battu, G. Wani, and A. A. Wani, ‘NER initiation factors, DDB2 and XPC, regulate UV radiation response by recruiting ATR and ATM kinases to DNA damage sites’, DNA Repair, vol. 12, no. 4, pp. 273–283, Apr. 2013, doi: 10.1016/j.dnarep.2013.01.003.

[72] J. E. Cleaver, R. R. Laposa, and C. L. Limoli, ‘DNA Replication in the Face of (In)surmountable Odds’, Cell Cycle, vol. 2, no. 4, pp. 309–314, Jul. 2003, doi: 10.4161/cc.2.4.436.

[73] L. Zannini, D. Delia, and G. Buscemi, ‘CHK2 kinase in the DNA damage response and beyond’, Journal of Molecular Cell Biology, vol. 6, no. 6, pp. 442–457, Dec. 2014, doi: 10.1093/jmcb/mju045.

[74] X. Bi, D. M. Slater, H. Ohmori, and C. Vaziri, ‘DNA Polymerase κ Is Specifically Required for Recovery from the Benzo[a]pyrene-Dihydrodiol Epoxide (BPDE)-induced S-phase Checkpoint’, Journal of Biological Chemistry, vol. 280, no. 23, pp. 22343–22355, Jun. 2005, doi: 10.1074/jbc.M501562200.

