## Supplementary material for "Human CSB-deficient iPSCs exhibit impaired DNA damage repair and stress responses following BPDE exposure in an early developmental model": Supplememtary Tables

### **Supplementary Table 1:** Primers Table

| Gene | Primer Forward | Primer reverse | Company |
|---|---|---|---|
| **ATM** | ACACTGCGCGTATAAGCCAATC | TTTTCAACCAGTTTTCCGTTACTTC | Eurofins |
| **ATR** | TGTTGGGCCCACTTTATGCAGC | TAGAGACGACCTGAGACGACGC | Eurofins |
| **BAX** | GAGCTGCAGAGGATGATTGCCG | CCCCAGTTGAAGTTGCCGTCAG | Eurofins |
| **BRCA1** | AGCCTGGCCAACATGGTGAAA | CTTGGCTCACTGCAACCTCCAC | Eurofins |
| **CHK1** | GGATCAGCTTTTCCCAGCCCAC | TTCTGTGAGGATCCTGGGGTGC | Eurofins |
| **CHK2** | ACGGAGTTCACAACACAGCAGC | TCTACTAGTCGAAAGCGGCCCC | Eurofins |
| **DDB2** | GCCATCTGTCCAGCA | GGGGTGAGTTGGGTC | Eurofins |
| **NANOG** | CCTGTGATTTGTGGGCCTG | GACAGTCTCCGTGTGAGGCAT | Eurofins |
| **OCT4** | AGTTTGTGCCAGGGTTTTTG | ACTTCACCTTCCCTCCAACC | Eurofins |
| **P53** | CAGGGCAGCTACGGTTTCC | CAGTTGGCAAAACATCTTGTTGAG | Eurofins |
| **RPLP0** | TCGACAATGGCAGCATCTAC | ATCCGTCTCCACAGACAAGG | Eurofins |
| **SOX2** | GTATCAGGAGTTGTCAAGGCAGAG | TCCTAGTCTTAAAGAGGCAGCAAAC | Eurofins |
| **XPC** | GACCTGAAGAAGGCACACCA | TGGCTTCACAGGCAGAAGAG | Eurofins |

### **Supplementary Table 2:** Antibodies Table

| Primary Ab | Species | Application | Dilution | company | Catalog number |
|---|---|---|---|---|---|
| **53BP1** | Mouse | IF | 1:200 | | |
| **BAX** | Rabbit | WB | 1:1000 | Cell Signaling Technology | 2772T |
| **BCL2** | Rabbit | WB | 1:1000 | Cell Signaling Technology | 4223S |
| **Cleaved Caspase 3** | Rabbit | WB | 1:1000 | Cell Signaling Technology | 9664S |
| **YH2AX** | Rabbit | WB/IF | 1:1000/1:200 | Cell Signaling Technology | 9718S |
| **CSB** | Rabbit | WB | 1:1000 | GeneTex | GTX104589 |
| **HSP90** | Mouse | WB | 1:1000 | BD Transduction Laboratories | 610419 |
| **NANOG** | Rabbit | WB/IF | 1:1000/1:200 | Cell Signaling Technology | 4903S |
| **OCT4A** | Rabbit | WB/IF | 1:1000/1:200 | Cell Signaling Technology | 2840S |
| **PAX6** | Rabbit | IF | 1:200 | Cell Signaling Technology | 60433S |
| **P-CHK2** | Rabbit | WB | 1:1000 | Cell Signaling Technology | 2661s |
| **P-SMAD 1/5** | Rabbit | WB | 1:1000 | Cell Signaling Technology | 9516S |
| **P-SMAD 2** | Rabbit | WB | 1:1000 | Cell Signaling Technology | 3108S |
| **Anti-p53 (Ab-1) (Pantropic) Mouse mAb (PAb421)** | Mouse | WB | 1:1000 | Millipore | OP03 |
| **SMAD 2/3** | Rabbit | WB | 1:1000 | Cell Signaling Technology | 8685S |
| **SMAD 1** | Rabbit | WB | 1:1000 | Cell Signaling Technology | 6944S |

| | | | | | |
|---|---|---|---|---|---|
| **SOX2** | Rabbit | WB/IF | 1:1000/1:200 | Cell Signaling Technology | 3579S |
| **SSEA4** | Mouse | IF | 1:200 | Cell Signaling Technology | 4755S |
| **TRA-1-60** | Mouse | IF | 1:200 | Cell Signaling Technology | 4746S |
| **Tra-1-81** | Mouse | IF | 1:200 | Cell Signaling Technology | 4745S |

| Secondary Ab | Application | Dilution | company | Catalog number |
|---|---|---|---|---|
| **Alexa 555 gt anti mouse IgG (H+L)** | IF | 1:400 | ThermoFisher | A21424 |
| **Alexa 488 dk anti rb IgG (H+L)** | IF | 1:400 | ThermoFisher | A21206 |
| **Alexa 488 gt anti rb IgG (H+L)** | IF | 1:400 | ThermoFisher | A11008 |
| **Alexa 488 dk anti gt IgG (H+L)** | IF | 1:400 | ThermoFisher | A11055 |
| **Alexa 555 gt Anti-Rb IgG(H+L)** | IF | 1:400 | ThermoFisher | A21428 |
| **Anti-Rabbit IgG, HRP linked Ab** | WB | 1:2500 | CST | 7074S |
| **ECL peroxidase laabelled anti Mouse Ab** | WB | 1:2500 | GE | NA931V |
