## Supplementary material for "Human CSB-deficient iPSCs exhibit impaired DNA damage repair and stress responses following BPDE exposure in an early developmental model": Supplememtary Figures

[§] Corresponding author

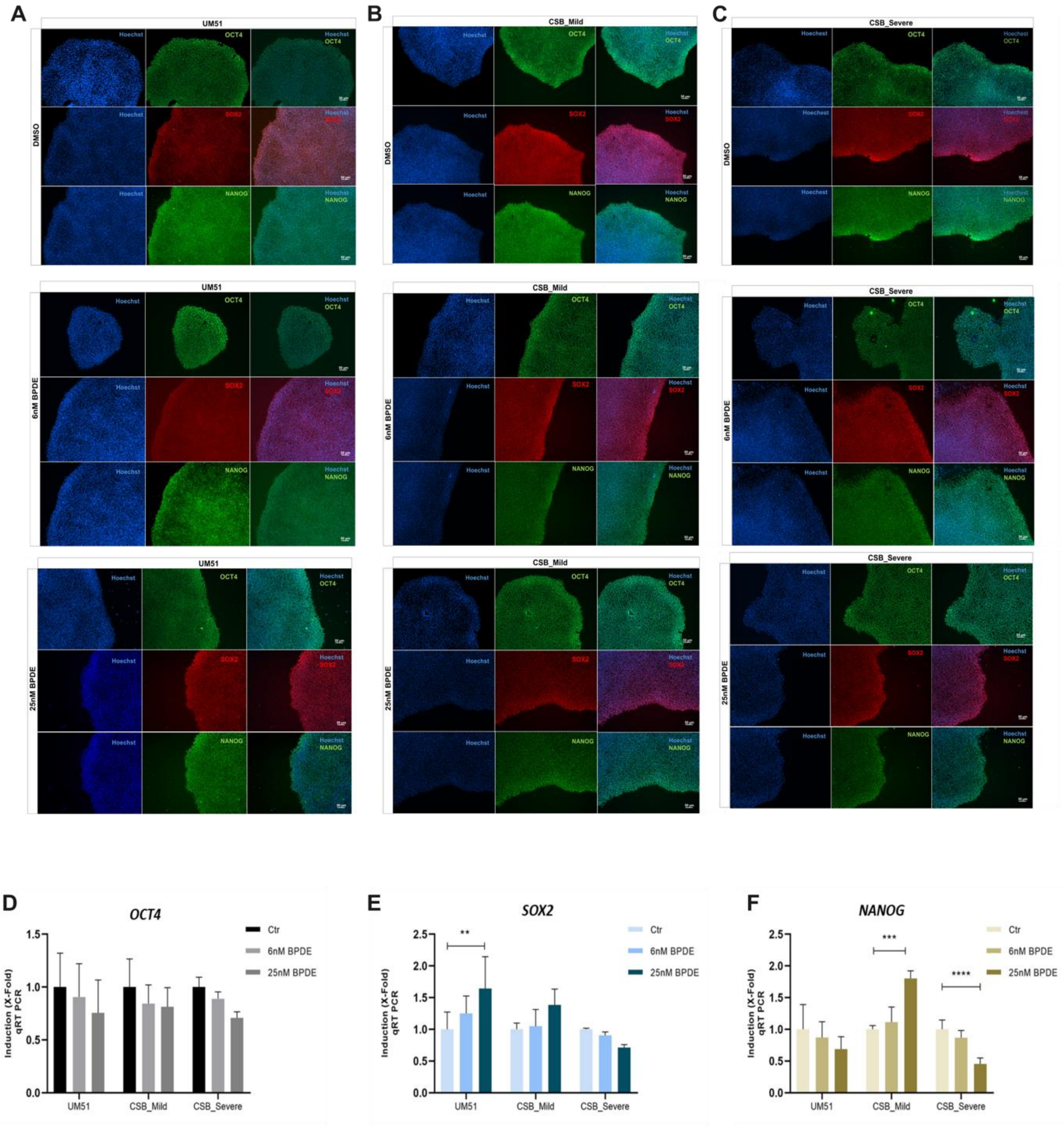


**Supplementary Figure 1: Pluripotency marker transcript levels in iPSCs following BPDE treatment.** iPSC cultures were treated with 6 nM or 25 nM BPDE for 24 hours prior to fixation or protein/RNA extraction. **(A-C)** Immunofluorescence staining of pluripotency markers OCT4 (green), SOX2 (red) and NANOG (green) in UM51 (A), CSB_Mild (B), and CSB_Severe (C) iPSCs following 24-hour exposure to 6 nM or 25 nM BPDE. Scale bar: 50 µm. (**D-F)** Quantitative RT-PCR analysis of pluripotency markers *OCT4* (D), *SOX2* (E), and *NANOG* (F) (OSN) in UM51, CSB_Mild, and CSB_Severe iPSCs following BPDE exposure. Expression values were normalized to *RPL0* and are presented as mean ± 95 % confidence interval ($n$ = 3). Statistical significance is indicated as *$p$ < 0.05, **p < 0.01, ***$p$ < 0.001 and ****$p$ < 0.0001.

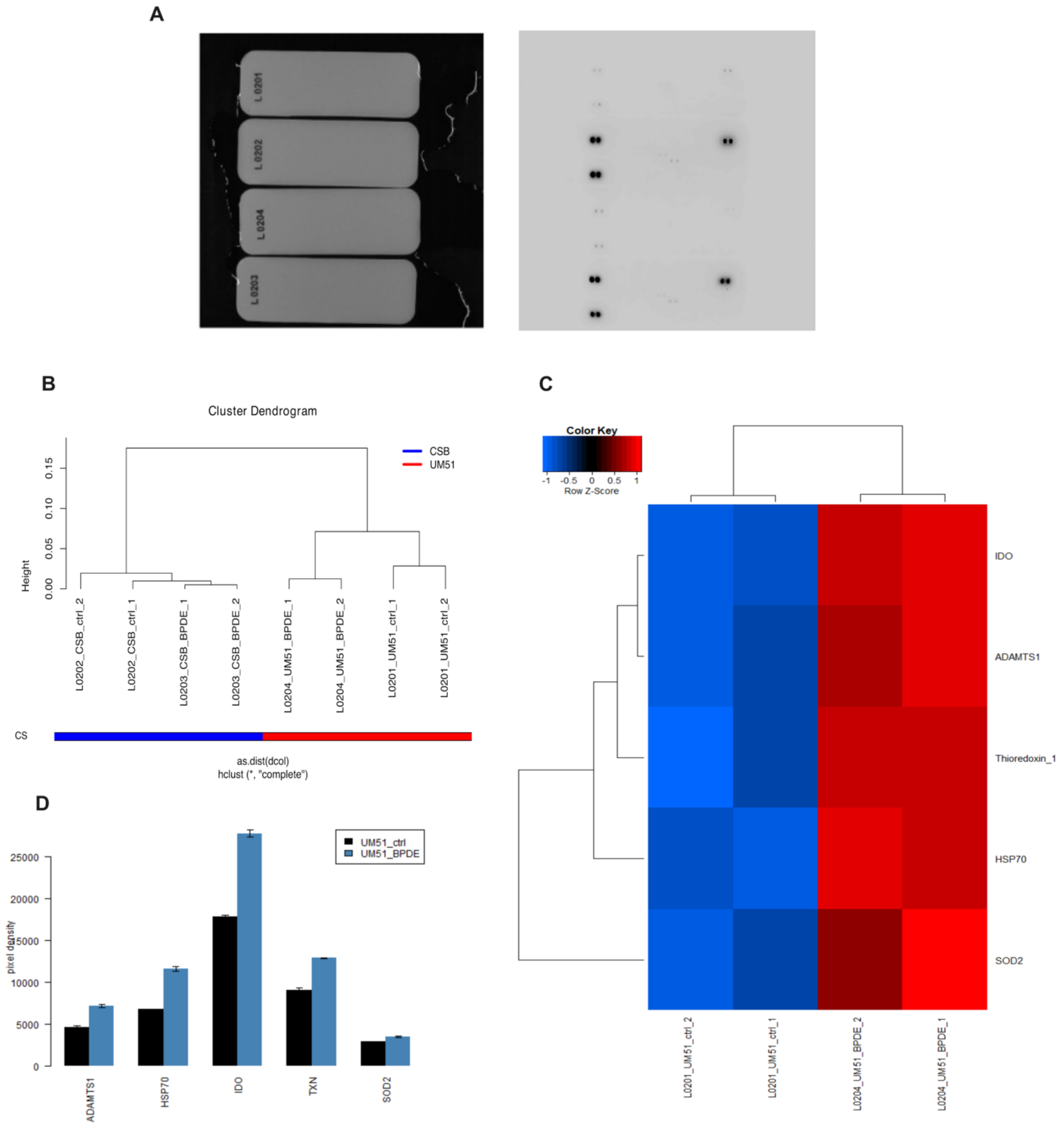


**Supplementary Figure 2: Stress array analysis of UM51 and CSB_Severe iPSCs following BPDE exposure.** iPSC cultures were treated with 25 nM BPDE for 24 hours prior to protein extraction. **(A)** Photograph of the stress array membrane showing the relative levels of selected human cell stress–related proteins prior to electrochemiluminescence (ECL) measurement. Samples: UM51 DMSO (L0201), UM51 25 nM BPDE (L0204), CSB_Severe DMSO (L0202), and CSB_Severe 25 nM BPDE (L0203). ECL signals were detected after 10 min, showing localized luminescence corresponding to protein expression at specific detection sites. **(B)** Cluster dendrogram illustrating the relationship between UM51 and CSB_Severe iPSCs following BPDE exposure. **(C)** Heatmaps depicting expression profiles of the most affected proteins in UM51 cells after 24-hour exposure to 25 nM BPDE. **(D)** Bar chart summarizing the expression levels of the most affected proteins in the stress array of UM51 cells following 24-hour exposure to 25 nM BPDE.

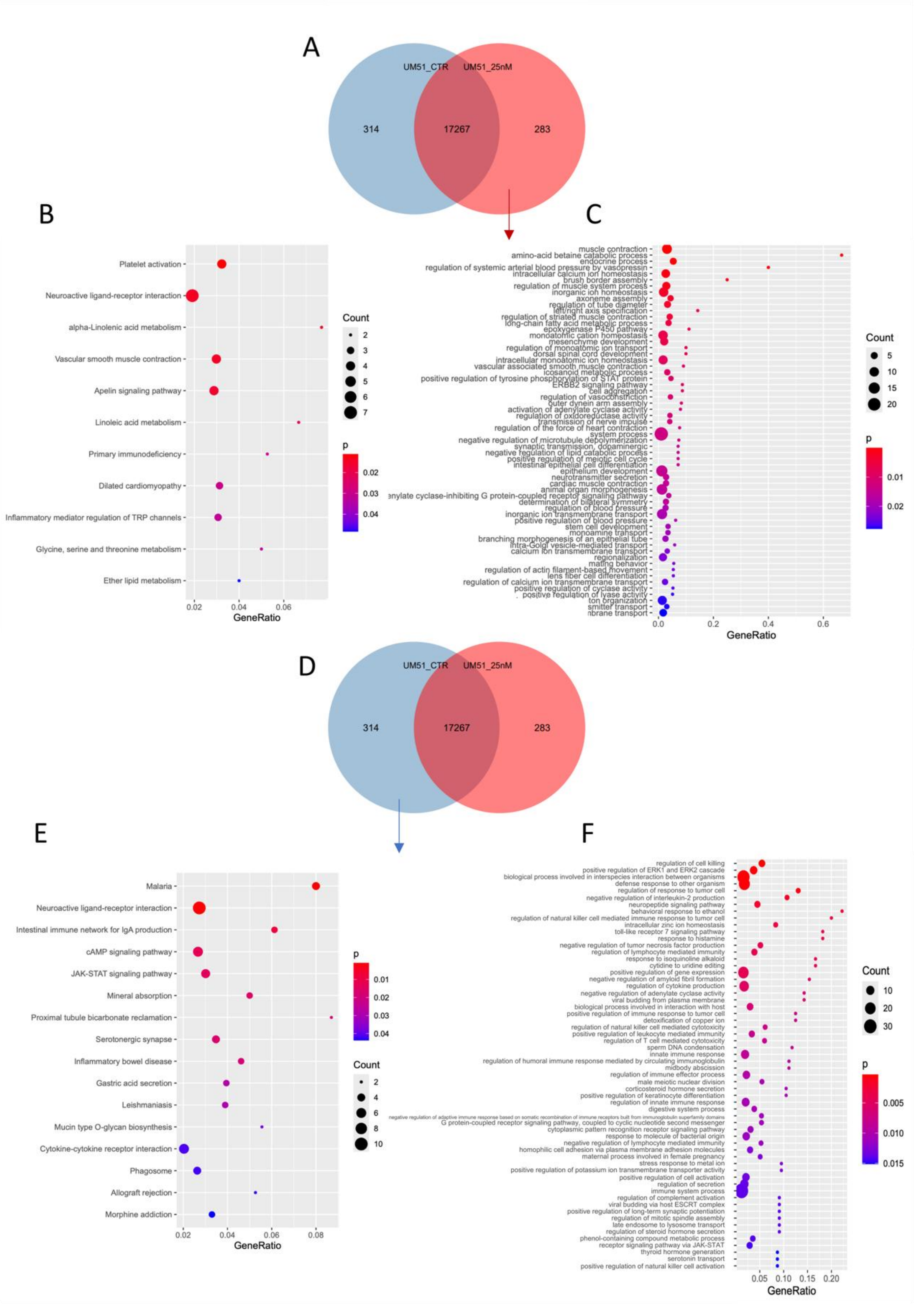
A
UM51_CTR
UM51_25nM
314
17267
283
B
Platelet activation
Neuroactive ligand-receptor interaction
alpha-Linolenic acid metabolism
Vascular smooth muscle contraction
Apelin signaling pathway
Linoleic acid metabolism
Primary immunodeficiency
Dilated cardiomyopathy
Inflammatory mediator regulation of TRP channels
Glycine, serine and threonine metabolism
Ether lipid metabolism
0.02
0.04
0.06
GeneRatio
Count
2
3
4
5
6
7
p
0.02
0.03
0.04
C
muscle contraction
amino-acid betaine catabolic process
endocrine process
regulation of systemic arterial blood pressure by vasopressin
intracellular calcium ion homeostasis
brush border assembly
regulation of muscle system process
inorganic ion homeostasis
axoneme assembly
regulation of tube diameter
left/right axis specification
regulation of striated muscle contraction
long-chain fatty acid metabolic process
epoxygenase P450 pathway
monoatomic cation homeostasis
mesenchyme development
regulation of monoatomic ion transport
dorsal spinal cord development
intracellular monoatomic ion homeostasis
vascular associated smooth muscle contraction
icosanoid metabolic process
positive regulation of tyrosine phosphorylation of STAT protein
ERBB2 signaling pathway
cell aggregation
regulation of vasoconstriction
outer dynein arm assembly
activation of adenylate cyclase activity
regulation of oxidoreductase activity
transmission of nerve impulse
regulation of the force of heart contraction
system process
negative regulation of microtubule depolymerization
synaptic transmission, dopaminergic
negative regulation of lipid catabolic process
positive regulation of meiotic cell cycle
intestinal epithelial cell differentiation
epithelium development
neurotransmitter secretion
cardiac muscle contraction
animal organ morphogenesis
enylate cyclase-inhibiting G protein-coupled receptor signaling pathway
determination of bilateral symmetry
regulation of blood pressure
inorganic ion transmembrane transport
positive regulation of blood pressure
stem cell development
monoamine transport
branching morphogenesis of an epithelial tube
intra-Golgi vesicle-mediated transport
calcium ion transmembrane transport
regionalization
mating behavior
regulation of actin filament-based movement
lens fiber cell differentiation
regulation of calcium ion transmembrane transport
positive regulation of cyclase activity
positive regulation of lyase activity
ton organization
smitter transport
nbrane transport
0.0
0.2
0.4
0.6
GeneRatio
Count
5
10
15
20
p
0.01
0.02
D
UM51_CTR
UM51_25nM
314
17267
283
E
Malaria
Neuroactive ligand-receptor interaction
Intestinal immune network for IgA production
cAMP signaling pathway
JAK-STAT signaling pathway
Mineral absorption
Proximal tubule bicarbonate reclamation
Serotonergic synapse
Inflammatory bowel disease
Gastric acid secretion
Leishmaniasis
Mucin type O-glycan biosynthesis
Cytokine-cytokine receptor interaction
Phagosome
Allograft rejection
Morphine addiction
0.02
0.04
0.06
0.08
GeneRatio
p
0.01
0.02
0.03
0.04
Count
2
4
6
8
10
F
regulation of cell killing
positive regulation of ERK1 and ERK2 cascade
biological process involved in interspecies interaction between organisms
defense response to other organism
regulation of response to tumor cell
negative regulation of interleukin-2 production
neuropeptide signaling pathway
behavioral response to ethanol
regulation of natural killer cell mediated immune response to tumor cell
intracellular zinc ion homeostasis
toll-like receptor 7 signaling pathway
response to histamine
negative regulation of tumor necrosis factor production
regulation of lymphocyte mediated immunity
response to isoquinoline alkaloid
cytidine to uridine editing
positive regulation of gene expression
negative regulation of amyloid fibril formation
regulation of cytokine production
negative regulation of adenylate cyclase activity
viral budding from plasma membrane
biological process involved in interaction with host
positive regulation of immune response to tumor cell
detoxification of copper ion
regulation of natural killer cell mediated cytotoxicity
positive regulation of leukocyte mediated immunity
regulation of T cell mediated cytotoxicity
sperm DNA condensation
innate immune response
regulation of humoral immune response mediated by circulating immunoglobulin
midbody abscission
regulation of immune effector process
male meiotic nuclear division
corticosteroid hormone secretion
positive regulation of keratinocyte differentiation
regulation of innate immune response
digestive system process
G protein-coupled receptor signaling pathway, coupled to cyclic nucleotide second messenger
cytoplasmic pattern recognition receptor signaling pathway
response to molecule of bacterial origin
negative regulation of lymphocyte mediated immunity
homophilic cell adhesion via plasma membrane adhesion molecules
maternal process involved in female pregnancy
stress response to metal ion
positive regulation of potassium ion transmembrane transporter activity
positive regulation of cell activation
regulation of secretion
immune system process
regulation of complement activation
viral budding via host ESCRT complex
positive regulation of long-term synaptic potentiation
regulation of mitotic spindle assembly
late endosome to lysosome transport
regulation of steroid hormone secretion
phenol-containing compound metabolic process
receptor signaling pathway via JAK-STAT
thyroid hormone generation
serotonin transport
positive regulation of natural killer cell activation
0.05
0.10
0.15
0.20
GeneRatio
Count
10
20
30
p
0.005
0.010
0.015

**Supplementary Figure 3: Global transcriptome and pathway analysis of UM51 iPSCs following BPDE exposure.**

iPSC cultures at approximately 90 % confluency were treated with 25 nM BPDE for 24 hours prior to RNA extraction. **(A)** Venn diagram showing genes uniquely expressed in control UM51 iPSCs, genes uniquely expressed in 25 nM BPDE–treated UM51 iPSCs, and genes common to both conditions (detection $p < 0.05$). **(B)** Dot plots depicting the top KEGG pathways enriched among genes uniquely expressed in BPDE-treated UM51 iPSCs. **(C)** Gene Ontology (GO) analysis of genes uniquely expressed in BPDE-treated UM51 iPSCs. **(D)** Venn diagram showing genes uniquely expressed in control UM51 iPSCs, genes uniquely expressed in BPDE-treated UM51 iPSCs, and genes common to both conditions (detection $p < 0.05$). **(E)** Dot plots depicting the top KEGG pathways enriched among genes uniquely expressed in control UM51 iPSCs. **(F)** Gene Ontology (GO) analysis of genes uniquely expressed in control UM51 iPSCs.

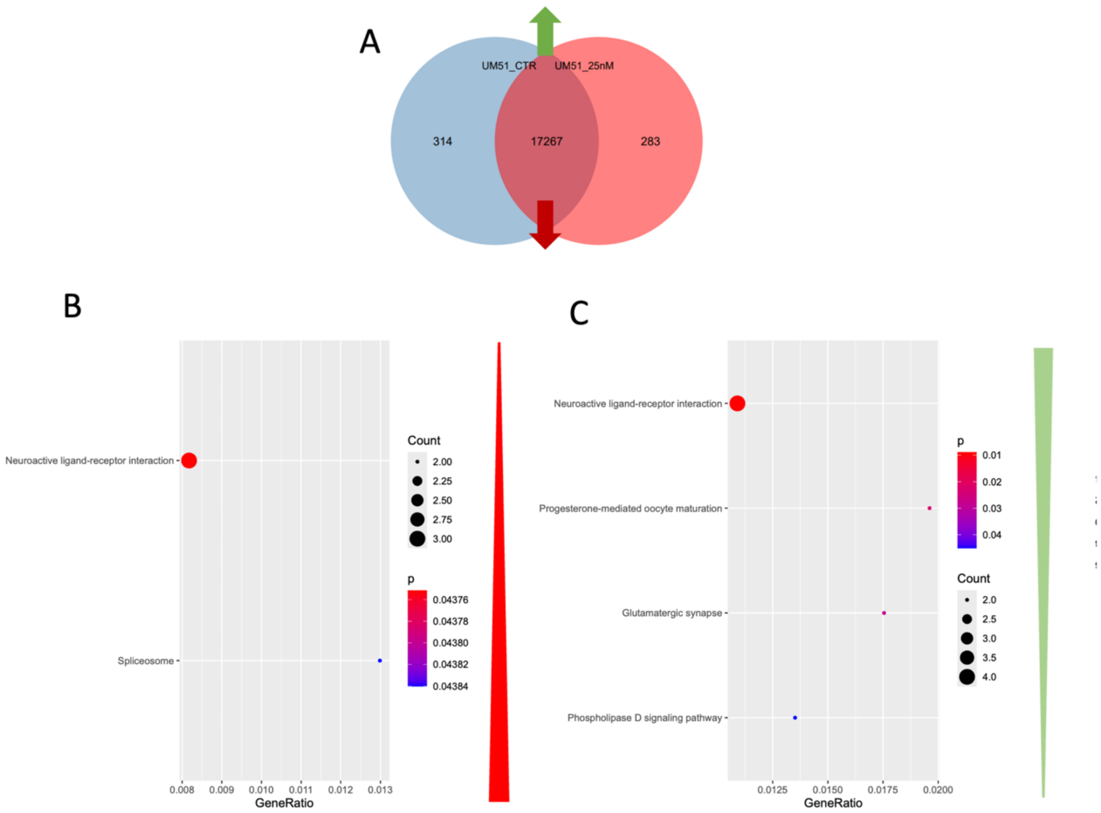


**Supplementary Figure 4. Global transcriptome and associated pathway analysis of UM51 iPSCs following BPDE exposure.**

iPSC cultures at approximately 90 % confluency were treated with 25 nM BPDE for 24 hours prior to RNA extraction. **(A)** Venn diagram showing genes uniquely expressed in control **UM51** iPSCs, genes uniquely expressed in 25 nM BPDE–treated UM51 iPSCs, and genes common to both conditions (detection $p < 0.05$). **(B)** Dot plots depicting the top upregulated KEGG pathways in UM51 iPSCs after genotoxic stress. **(C)** Dot plots depicting the top downregulated KEGG pathways in UM51 iPSCs after genotoxic stress.

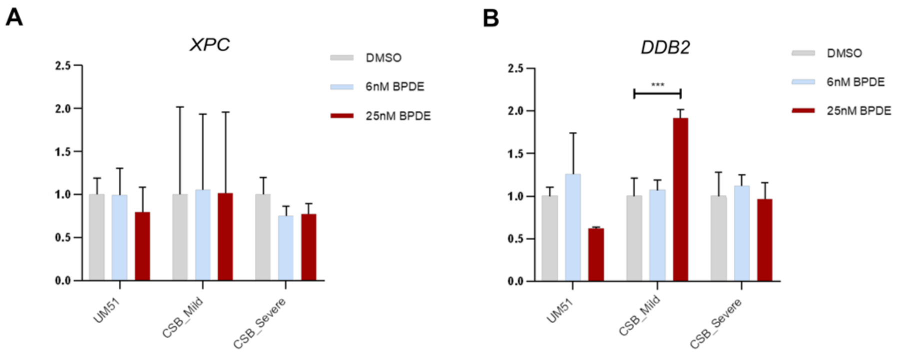


**Supplementary Figure 5: Transcriptional analysis of DNA repair genes in iPSCs following BPDE exposure.**
**(A)** Quantitative RT-PCR analysis of *XPC* transcript levels in iPSCs after 24 hours of BPDE treatment. **(B)** Quantitative RT-PCR analysis of *DDB2* transcript levels in iPSCs after BPDE treatment. Expression values were normalized to *RPLP0* and are presented as mean ± 95 % confidence interval ($n$ = 3). Statistical significance is indicated as *$p < 0.05$, **p < 0.01, ***$p < 0.001$ and ****$p < 0.0001$.

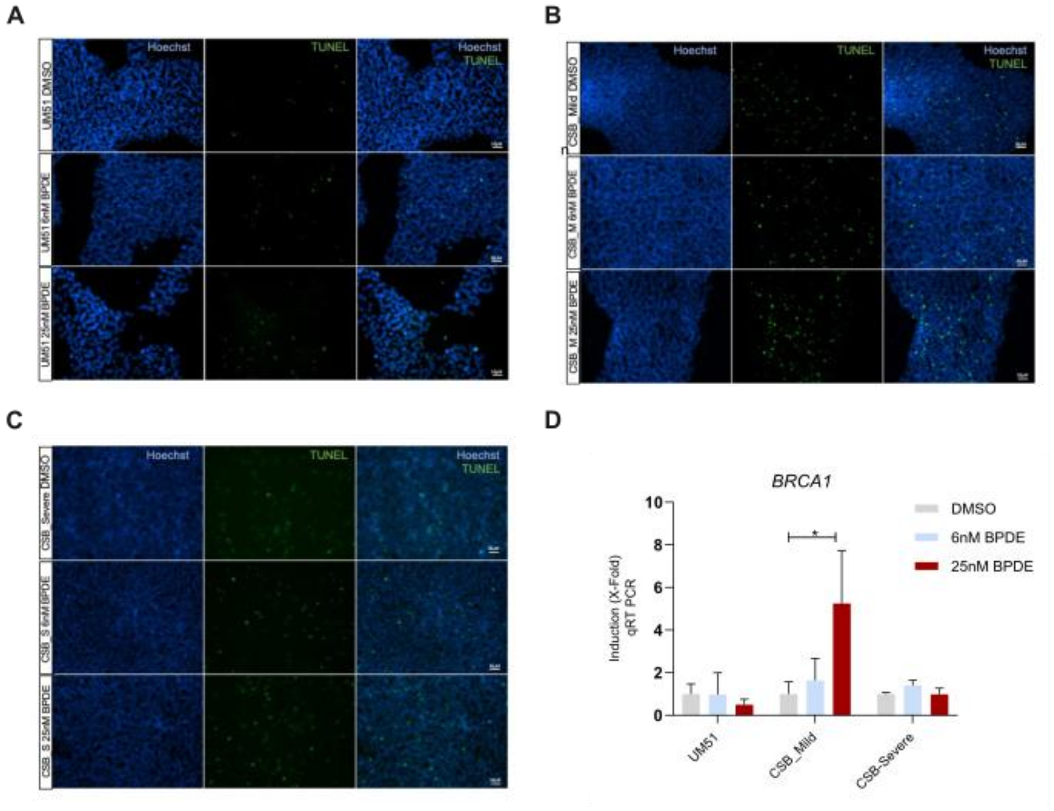


**Supplementary Figure 6:**
iPSC cultures were treated with 6 nM or 25 nM BPDE for 24 hours prior to fixation or RNA extraction. **(A-C)** Immunofluorescence staining of the apoptotic marker TUNEL (green) and Hoechst (blue) in UM51 (A), CSB_Mild (B), and CSB_Severe (C) iPSCs following 24-hour exposure to 6 nM or 25 nM BPDE. Scale bar: 50 µm.
